# Exploring neural tracking of acoustic and linguistic speech representations in individuals with post-stroke aphasia

**DOI:** 10.1101/2023.03.01.530707

**Authors:** Jill Kries, Pieter De Clercq, Marlies Gillis, Jonas Vanthornhout, Robin Lemmens, Tom Francart, Maaike Vandermosten

**Affiliations:** Experimental Oto-Rhino-Laryngology, Department of Neurosciences, Leuven Brain Institute, KU Leuven, Leuven, Belgium; Department of Psychology, Stanford University, Stanford, CA, USA; Experimental Neurology, Department of Neurosciences, KU Leuven, Leuven, Belgium; Laboratory of Neurobiology, VIB-KU Leuven Center for Brain and Disease Research, Leuven, Belgium; University Hospitals Leuven, Department of Neurology, Leuven, Belgium

**Keywords:** speech processing, neural tracking, aphasia, EEG, stroke

## Abstract

Aphasia is a communication disorder that affects processing of language at different levels (e.g., acoustic, phonological, semantic). Recording brain activity via EEG while people listen to a continuous story allows to analyze brain responses to acoustic and linguistic properties of speech. When the neural activity aligns with these speech properties, it is referred to as neural tracking. Even though measuring neural tracking of speech may present an interesting approach to studying aphasia in an ecologically valid way, it has not yet been investigated in individuals with stroke-induced aphasia. Here, we explored processing of acoustic and linguistic speech representations in individuals with aphasia in the chronic phase after stroke and age-matched healthy controls. We found decreased neural tracking of acoustic speech representations (envelope and envelope onsets) in individuals with aphasia. In addition, word surprisal displayed decreased amplitudes in individuals with aphasia around 195 ms over frontal electrodes, although this effect was not corrected for multiple comparisons. These results show that there is potential to capture language processing impairments in individuals with aphasia by measuring neural tracking of continuous speech. However, more research is needed to validate these results. Nonetheless, this exploratory study shows that neural tracking of naturalistic, continuous speech presents a powerful approach to studying aphasia.

**Key points:**

- Individuals with aphasia display decreased encoding of acoustic speech properties (envelope and its onsets) in comparison to healthy controls.
- Neural responses to word surprisal reveal decreased amplitudes in individuals with aphasia around 195 ms processing time (not corrected for multiple comparisons).
- Neural tracking of natural speech can be used to study speech processing impairments in aphasia.

## 1 Introduction

About one third of strokes result in aphasia, a language disorder that can impact auditory comprehension, oral production, writing and/or reading (Engelter et al., 2006; National Aphasia Association, 2022; Pasley and Knight, 2013). Aphasia can impact communication to different degrees, ranging from subtle to severe impairments, and from recovery within hours after stroke to permanent language impairments. The severity and persistence depend on factors such as lesion location and size, brain plasticity, therapy, intrinsic motivation and social support (Pasley and Knight, 2013; Schevenels et al., 2020; Cordella et al., 2022; Schevenels et al., 2022). In order to be effective, speech therapy should be given with high intensity and the content should be tailored to the specific problems of each individual with aphasia (IWA) (Rohde et al., 2018; Engelter et al., 2006; Schevenels et al., 2020; Brady, 2022). Individually tailored therapy requires a precise diagnosis of language impairments.

Although diagnostic tests originally focused on the classic aphasia typology (i.e., assessing performance on fluency, comprehension and repetition tasks), neuroimaging studies have suggested that focusing on the different language processing components (i.e., acoustic, phonological, semantic, syntactic) is more in line with the neural networks of language processing (e.g., the dual-stream model by Hickok and Poeppel (2007)) (Rohde et al., 2018; Tremblay and Dick, 2016; Pasley and Knight, 2013; Wilson et al., 2023). Fridriksson et al. (2018) suggest that lesions in the dorsal stream (i.e., sensori-motor integration) may impair phonological processing, whereas damage to the ventral stream (i.e., acoustic-semantic integration) may impair semantic processing. Hence, a neuroimaging approach focused on the linguistic aspects may yield more precise diagnostic insights and a more effective therapeutic approach (Tremblay and Dick, 2016; Pasley and Knight, 2013). Moreover, behavioral testing after stroke is difficult because in 80% of cases, IWA one year post-stroke have co-morbid cognitive problems, such as memory, executive functions and/or attention problems, which can bias the results or even impede behavioral testing (El Hachioui et al., 2014; Fonseca et al., 2018). Further, behavioral tests consist of artificial tasks that do not always correspond to communication abilities in daily life. Thus, to provide targeted intervention and improve recovery outcomes, an aphasia diagnosis that provides precise insights, that is less dependent on cognitive performance and that is more ecologically valid is needed.

Electroencephalography (EEG) provides ways to study the brain’s responses to speech with reduced active participation of the patient. By averaging the EEG signal in response to a large number of repetitive sound or speech stimuli, peaks in specific time ranges have been consistently identified in neurotypicals, i.e., event-related potentials (ERPs), offering a window into the spatio-temporal patterns of the neural response to speech. This way, ERPs related to acoustic (e.g., P1-N1-P2 complex (e.g., Martin et al., 2008; Harris, 2020)) and linguistic (e.g., N400 (e.g., Hillyard and Kutas, 1984; Kutas and Federmeier, 2011; Nieuwland et al., 2020)) aspects of speech have been identified. In IWA, altered ERPs have been found across language processing levels and across a variety of experimental stimuli and tasks (Ofek et al., 2013; Becker and Reinvang, 2007; Ilvonen et al., 2001; Pulvermüller et al., 2004; Aerts et al., 2015; Ilvonen et al., 2004; Pettigrew et al., 2005; Robson et al., 2017; Chang et al., 2016; Kawohl et al., 2010; Khachatryan et al., 2017; Sheppard et al., 2017; Lice and Palmović, 2017; Kielar et al., 2012; Räling et al., 2016). Most of these studies have reported decreased amplitudes and increased latencies in IWA as compared to healthy controls, with the exception of the P2 in Aerts et al. (2015) and Ilvonen et al. (2001), which observed opposite patterns. Some of these studies have found ERP amplitudes or latencies to be correlated with language performance (Pettigrew et al., 2005; Robson et al., 2017; Khachatryan et al., 2017).

The potential of ERPs to serve as evaluatory measure of intervention effects has recently been reviewed by Cocquyt et al. (2020), who concluded that there is potential for ERPs to assess levels of aphasia symptoms, after development of normative data. However, to date ERPs are not commonly used in the clinic. This is likely due to small sample sizes and the heterogeneity of aphasia symptoms within the studied samples (Silkes and Anjum, 2021), which complicates the development of validated norms for ERPs. While ERPs are useful to understand the functional meaning of different peaks in the spatio-temporal patterns of the neural response, their application in aphasia diagnostics poses further challenges, e.g., long administration time due to different paradigms at distinct speech processing levels and the need for a large number of repetitive stimuli to average across (Kandylaki and Bornkessel-Schlesewsky, 2019). Moreover, listening to repetitive and artificially created stimuli is not representative of everyday language situations that IWA struggle with mostly (Le et al., 2018). More naturalistic speech stimuli, such as a narrative, would present a more ecologically valid stimulus to analyze the brain’s response to speech (Lalor and Foxe, 2010; Ding and Simon, 2012; Hamilton and Huth, 2018; Kandylaki and Bornkessel-Schlesewsky, 2019; Gillis et al., 2022).

From the narrative, different characteristics or representations of speech can be derived and their relation to the EEG signal can be measured. When the neural signals align with the speech properties, it is referred to as neural tracking. By examining the data in this way, spatial and temporal neural response properties in response to multiple speech representation levels (e.g., acoustic, phonological, semantic, syntactic) can be analyzed from the same data (Di Liberto et al., 2015; Brodbeck et al., 2018; Gillis et al., 2021, 2022; Mesik et al., 2021). Moreover, a measure of the strength with which the different speech representations are encoded in the EEG signal can be computed. Research has shown that even relatively short EEG recordings (i.e., 10-20 minutes) can provide valid results with this approach (Di Liberto and Lalor, 2017). Furthermore, limited active participation is required from the participant during such a paradigm, which is especially advantageous for testing IWA. These characteristics make neural tracking an ideal tool to study aphasia.

Examining neural tracking, the most frequently studied speech representation to date is the speech envelope, consisting of the slow amplitude modulations of speech over time (Aiken and Picton, 2008). The envelope presents an essential cue for speech intelligibility (Aiken and Picton, 2008; Shannon et al., 1995). Whereas envelope tracking has not yet been investigated in individuals with stroke-induced aphasia, Dial et al. (2021) have explored it in individuals with primary progressive aphasia (PPA). Individuals with the logopenic variant of this neurodegenerative disease displayed increased envelope tracking compared to healthy controls in the theta band (Dial et al., 2021). On the other hand, envelope tracking in other disorders, such as developmental dyslexia, have shown decreased envelope tracking compared to controls in the 1-8 Hz range, though the results were largely driven by the delta band (Di Liberto et al., 2018). These studies show that neural tracking of continuous speech presents a promising new avenue to study language disorders.

While envelope tracking is mostly considered an acoustic process in the literature, it has been found to be affected by speech intelligibility and higher-level speech-specific processes (Peelle et al., 2013; Vanthornhout et al., 2018; Broderick et al., 2019; Prinsloo and Lalor, 2022). This is not surprising given that the speech envelope also encompasses important cues for segmentation (i.e., rise and fall times to identify acoustic edges) of the continuous speech signal into discrete units (i.e., phonemes, syllables, words, phrases) (Aiken and Picton, 2008). Moreover, the syllable stress is also comprised in the envelope, which gives the listener an indication of the prosody and thus even conveys linguistic information. Given this entanglement of acoustic and linguistic cues, the speech envelope alone may not be ideal to find specific neurophysiological correlates of acoustic, phonological and lexical semantic processing in aphasia.

Recently, neural tracking of higher-level linguistic speech representations has been investigated, i.e., speech representations containing information about phonemes and words that take into account linguistic context (Broderick et al., 2018; Brodbeck et al., 2018; Weissbart et al., 2019; Gillis et al., 2021). This way, it has been observed that older adults, in comparison to younger adults, have an altered neural response to linguistic processes (Broderick et al., 2021; Mesik et al., 2021; Gillis and Kries et al., 2023). To date, higher-level linguistic speech representations have not been investigated in IWA via neural tracking, although higher-level speech processing impairments at the phonological and semantic level are reported most frequently in IWA (in contrast to acoustic processing impairments).

In the present study, we investigated neural tracking of acoustic and linguistic speech representations in individuals with post-stroke aphasia and healthy, age-matched controls. To this end, EEG data was acquired while participants listened to a continuous story, of which 8 speech representations were derived. The envelope and envelope onsets were considered to be acoustic representations of speech. The phoneme and word onsets were considered representations of speech segmentation, i.e., at the interface between acoustic and linguistic information. Lastly, phoneme surprisal and phoneme entropy were considered to be linguistic representations at the phoneme level, while word surprisal and word frequency were regarded as linguistic representations at the word level, i.e., related to lexical meaning. For the linguistic speech representations, we controlled the variance explained by acoustic cues and vice-versa, as we aimed to disentangle different levels of speech processing. Ultimately, disentangling mechanisms at different levels of speech processing is necessary to investigate whether neural tracking will be useful as a diagnostic tool for aphasia that provides information about different speech processing aspects. Here, our aim as a first step towards this goal was to explore group differences between IWA and healthy controls based on neural tracking of the different speech representations. Specifically, we studied group differences regarding the strength of neural tracking, i.e., *how well* the brain tracks specific aspects of speech, and regarding *how* the spatio-temporal pattern of the neural response operates during continuous speech perception. Based on the aforementioned ERP and neural tracking studies, we expected to observe group differences in speech representations at both acoustic and linguistic levels.

## 2 Materials & methods

### 2.1 Participants

We tested 41 IWA in the chronic phase after stroke (*≥* 6 months) and 24 healthy controls that were age-matched at group-level. 2 IWA had to be excluded post hoc-one because no lesion could be found in the left hemisphere and one because we did not have access to any lesion information-resulting in a sample size of 39 IWA. IWA were recruited in two ways. Between October 2018 and April 2022 (with a COVID-19-related break between March and June 2020), patients were recruited via daily screening at the stroke unit of the university hospital Leuven (score *≤* cut-off threshold on the Language Screening Test (LAST) (Flamand-Roze et al., 2011)) or via advertising the study in speech-language pathologists’ practices and rehabilitation centra (patients with a formal aphasia diagnosis) (see supplementary fig. S.1 for a detailed flowchart). Healthy age-matched controls were recruited via flyers positioned in recreational community centers for elderly. The target age of healthy controls was gradually adapted based on the mean age of IWA included in the study (supplementary figure S.2). The participants in this study are partly overlapping with participants in Kries et al. (2023).

For data collection, we only included IWA that had no formal diagnosis of a psychiatric or neurodegenerative disorder and that had a left-hemispheric or bilateral lesion. All aphasia participants were tested in the chronic phase after stroke (time since stroke onset in months (median(range)): 16.1(6-126.1). The aphasia sample was checked for language impairments at the moment of data collection using two standardized diagnostic aphasia tests, i.e., the diagnostic test ScreeLing Visch-Brink et al. (2010) and the Dutch picture-naming test (Nederlandse Benoemtest (NBT); Van Ewijk et al. (2020), using the same procedure as reported in Kries et al. (2022, biorxiv). The ScreeLing was administered on a tablet using the Gorilla Experiment Builder (http://www.gorilla.sc) (Anwyl-Irvine et al., 2020). The average score by group is reported in table 1. We included individuals that scored either (1) below the cut-off threshold (ScreeLing threshold: 68/72 points; NBT threshold: 255/276 points) on at least one of these two tests at the moment of data collection (n=27) (supplementary table S.1; supplementary fig. S.3), or (2) had a documented language impairment in the acute phase (n=12). Note that 10 out of the latter 12 IWA still followed speech-language therapy at the time of data collection (supplementary table S.1).

**Table 1:** Demographics, language-diagnostic information and covariates by group.

|  |  | Control | Aphasia |  |
| --- | --- | --- | --- | --- |
| **Demographics** |  |  |  |  |
| Age in years (mean(SD))<sup>◊</sup> |  | 71.5(7) | 69.5(12.4) | n.s. |
| Sex (n(%)) | female | 8(33.3%) | 13(33.3%) | n.s. |
|  | male | 16(66.7%) | 26(66.7%) |  |
| Education (n(%)) | primary education | 0(0%) | 3(7.7%) | n.s. |
|  | secondary education | 5(20.8%) | 15(38.4%) |  |
|  | tertiary education/college | 8(33.3%) | 10(25.6%) |  |
|  | Master's degree | 9(37.5%) | 11(28.2%) |  |
|  | PhD | 2(8.3%) | 0(0%) |  |
| Handedness (n(%)) | right | 22(91.6%) | 35(89.7%) | n.s. |
|  | left | 1(4.2%) | 2(5.1%) |  |
|  | ambidextrous | 1(4.2%) | 2(5.1%) |  |
| Multilingual (n(%)) | yes | 22(91.7%) | 33(84.6%) | n.s. |
|  | no | 2(8.3%) | 6(15.3%) |  |
| **Language diagnostic tests** |  |  |  |  |
| ScreeLing (max=72) (mean(SD)) |  | 69.9(2.5) | 61.5(9.4) | *** |
| Picture naming (max=276) (mean(SD)) |  | 271.3(4.1) | 224.7(58.1) | *** |
| **Covariates** |  |  |  |  |
| Hearing (Fletcher index in dB HL) (mean(SD)) |  | 24.5(12) | 28.4(14.1) | n.s. |
| Cognition (OCS composite score in %) (mean(SD)) |  | 94.2(4.7) | 80.4(16.6) | *** |
| Alertness (Likert scale) (median(range)) |  | 4.75(3.5-5) | 3.5(2-5) | ** |
| Fatigue (Likert scale) (median(range)) |  | 1.5(1-4.5) | 3.5(1-4.5) | ** |

<sup>◊</sup> Age is also used as a covariate; n.s. = not significantly different; $p < 0.05^*$ ; $p < 0.01^{**}$ ; $p < 0.001^{***}$

All participants were Dutch native speakers from Flanders, Belgium. Informed consent was obtained from all participants for the recruitment via screening and for the data collection in the chronic phase. The study received ethical approval by the medical ethical committee of KU Leuven and UZ Leuven (S60007) and is in accordance with the declaration of Helsinki.

In table 1, we summarized demographic information by group (details can be found in supplementary table S.1). Age, sex, education, handedness and multilinguality did not differ between groups (age: W = 464, p = 0.96; sex: *χ*^2^ = 0, df = 1, p = 1; education: *χ*^2^ = 7.26, df = 4, p = 0.1; handedness: *χ*^2^ = 0.063, df = 2, p = 0.98; multilingual: *χ*^2^ = 0.182, df = 1, p = 0.66). Details about the stroke in IWA, i.e., time since stroke onset, stroke type, occluded blood vessel, lesion location and speech-language therapy, can be found in supplementary table S.1. To visualize the damaged brain tissue of IWA, a lesion overlap image was created (fig. 1). More information on the lesion delineation process can be found in the supplementary material S.1.1.4. Demographic information was acquired via a self-reported questionnaire. Handedness was assessed via the Edinburgh Handedness Inventory (Oldfield, 1971).

**Figure 1:**
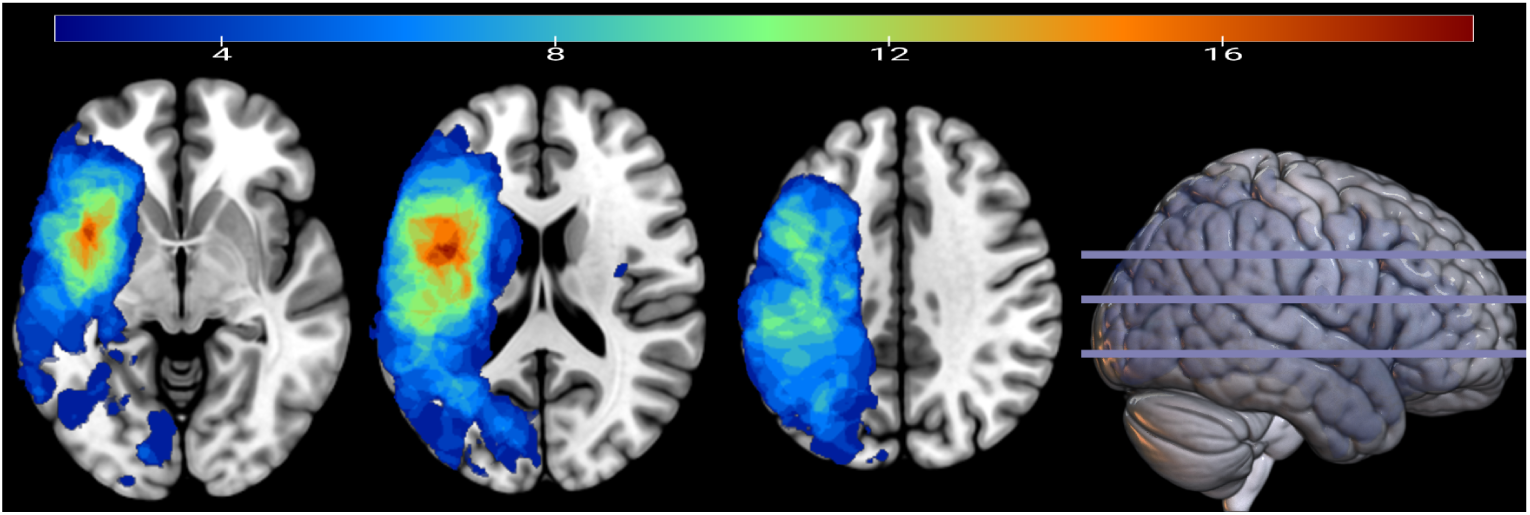
Lesion overlap image of the aphasia sample. The maximum overlap corresponds to 19 out of the total sample of 39 individuals with aphasia. Axial slices are shown in neurological orientation.

### 2.2 Behavioral measures that serve as covariates

#### 2.2.1 Hearing

Hearing thresholds were assessed via pure tone audiometry (PTA) at frequencies ranging from 0.25 to 4 kHz. In case the hearing thresholds below 4 kHz were *>*25 dB HL, this information was used to increase the amplitude of stimulus presentation during the EEG measurement. The PTA thresholds at 0.25, 0.5 and 1 kHz were averaged and then divided in half to come to the amount of dB that was added to the stimulus presentation amplitude of 60 dB SPL during the EEG paradigm. This calculation was done for each ear separately. After a short example stimulus, participants were asked whether the loudness was comfortable and if necessary the presentation volume was adjusted. The degree of volume adjustment did not differ between groups (W = 170, p = 0.8). Furthermore, hearing thresholds were used as covariates in statistical models (section 2.4). For this purpose, the Fletcher index (average of hearing thresholds at 0.5, 1 and 2 kHz) was calculated per ear and subsequently averaged across both ears. The Fletcher index did not differ between IWA and healthy controls (W = 541.5, p = 0.29, confidence interval: [-2.49 10]; table 1).

#### 2.2.2 Cognition

The Oxford Cognitive Screen-NL was administered to assess cognitive functioning (Huygelier et al., 2019). This test was designed to be language-independent, such that cognitive functioning can be disentangled from language functioning, which is especially important for IWA. Due to limited time in the testing protocol, we chose to only assess 4/10 subscales, i.e. attention and hemispatial neglect, reading, executive functioning, and memory. Hemispatial neglect was used as a means to potentially exclude participants in case they had too severe hemineglect, which could bias outcomes at most of the administered tests. However, the highest hemineglect score was still at a very mild level and thus we decided to not exclude any participants based on hemispatial neglect.

The task to assess attention consisted of crossing out target shapes among distractor shapes. The task to assess executive functions consisted of connecting circles and triangles in alternation in descending order of size. The memory task consisted of free recall and recognition of words (from the sentence read for the reading task) and shapes. These 3 tasks were used to calculate a composite score of cognitive functioning. This score was calculated by transforming the raw scores of each test into percentages and then averaging across the three outcomes. The composite score was used to regress out differences in cognitive functioning to explore neural tracking differences between groups. The cognition composite score was significantly lower in IWA than in healthy controls (W = 209.5, p *<* 0.001).

#### 2.2.3 Alertness and fatigue

Given that the experimental protocol (behavioral and EEG testing) was relatively long, especially considering that IWA often have cognitive impairments (e.g., attention), we decided to monitor the alertness and tiredness or fatigue at 3 time points throughout the experimental protocol (referred to as t1, t2 and t3). In supplementary figure S.4 E, the experimental protocol with reference to the timing of the alertness and fatigue questions is visualized. t1 was administered right at the start of the testing session, t2 after the EEG measurement and t3 at the end of the experimental protocol. Participants had to indicate on a Likert scale of 1 to 5 how alert and how tired they were at that moment. The questions were presented visually and auditory at the same time, as visualized in supplementary figure S.4 A and B. In supplementary section S.4, we describe the results of the interaction analysis between group and time points (supplementary fig. S.4 C and D, section S.1.1.5). Given that neural tracking has been shown to be influenced by attention (Lesenfants and Francart, 2020), we used the average ratings of t1 and t2 of the alertness and fatigue scale respectively as covariates in the analysis concerning group differences in neural tracking of speech. The average of t1 and t2 scores was specifically used because the EEG measurement took place in between these moments. A group difference was found for alertness (W = 243, p = 0.002) and for fatigue (W = 209, p = 0.001). IWA were on average less alert and more tired than healthy controls at t1 and t2 combined.

### 2.3 EEG-based measures

#### 2.3.1 Experimental paradigm

The EEG measurements took place in a soundproof room with Faraday cage. We recorded 64-channel EEG (ActiveTwo, BioSemi, Amsterdam, NL) at a sampling frequency of 8192 Hz. Participants were instructed to listen to a 24 minute long story while EEG data was recorded. They were seated one meter away from a screen and were asked to look at a fixation cross while listening in order to minimize eye movement artifacts in the EEG signal (fig. 2 A). The story *De wilde zwanen* (The Wild Swans), written by Hans Christian Andersen and narrated by a female Flemish-native speaker, was cut into 5 parts of on average 4.84 minutes (standard deviation (SD): 9.58 seconds) each. The silences in the story were reduced to 200 ms duration and the sample rate was set to 48 kHz. The software APEX (Francart et al., 2008) was used to calibrate and present stimuli. The story was presented bilaterally via shielded ER-3A insert earphones (Etymotic Research) at an amplitude of 60 dB SPL (A weighted), except if hearing thresholds were above 25 dB HL at the PTA, in which case the presentation volume was augmented (see section 2.2.1).

**Figure 2:**
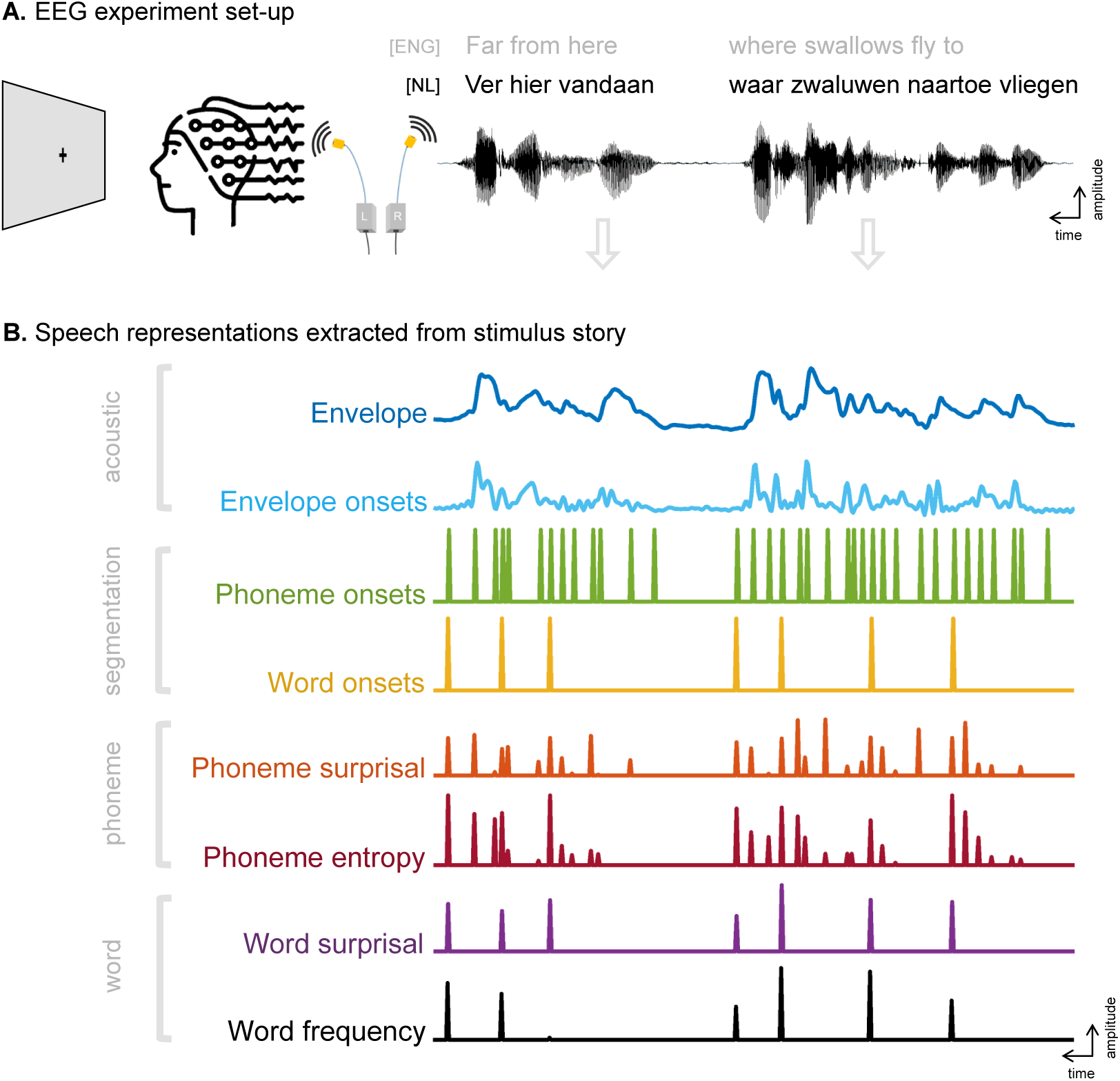
EEG experiment set-up and extraction of speech representations from the stimulus story. **A**. Participants listened to a story in Dutch (Flemish dialect) while EEG data was recorded. They were asked to look at a fixation cross while listening. The first 2 phrases of the story are visualized as written text (with an English translation) and audio signal. **B.** From the audio signal as depicted in panel A, 8 speech representations were extracted that reflect acoustic and linguistic properties of the story.

After each story part, participants answered a yes/no question and a multiple choice question about the content of the preceding story part. As these questions were not validated, we did not assess them. They were solely introduced in the protocol to make participants follow the content of the story attentively. Nonetheless, 5 participants (4 IWA, 1 control) fell asleep during parts of the EEG measurement. Given that an awake state is necessary to follow the contents of a story (such as reflected in linguistic speech representations) (Makov et al., 2017), we decided to exclude these story parts from the analysis. For one other control participant, a part of the data was not saved correctly and could thus also not be used for analysis. This means that for 6 participants, less than 24 minutes of data was used for analysis (19.36 minutes of data for 1 control and 1 aphasia participant, 14.52 minutes for 1 control and 2 aphasia participants, 9.68 minutes for 1 aphasia participant).

#### 2.3.2 EEG signal processing

The EEG signal processing was performed in MATLAB (version 9.1.0.441655 (R2016b)). The EEG data of the 5 story parts (i.e., epochs) were concatenated. Eye movement artifact removal was implemented using a multichannel Wiener filter (Somers et al., 2018). The EEG signal was referenced to the common average. For high-pass filtering, a least squares filter was applied with a filter order of 2000, with a passband frequency of 0.5 Hz and a stopband frequency of 0.45 Hz. For low-pass filtering, a least squares filter with a filter order of 2000 was applied with a passband frequency of 25 Hz and a stopband frequency of 27.5 Hz. The EEG data was downsampled to 128 Hz and subsequently normalized by subtracting the mean and dividing by the SD per epoch.

#### 2.3.3 Neural tracking

To investigate neural tracking, we used a forward modeling approach (i.e., encoding model), meaning that speech representations were used to predict the EEG signal (Mesgarani et al., 2014; Di Liberto et al., 2015; Holdgraf et al., 2017). Here, we were interested in both acoustic and linguistic speech representations. We relied on 8 representations, which were extracted from the stimulus story (fig. 2 B). We considered the envelope and envelope onsets as acoustic speech representations. Phoneme and word onsets represented phoneme– and word-level segmentation of speech. Phoneme surprisal and phoneme entropy were considered as linguistic representations at the phoneme level, word surprisal and word frequency at the word level.

##### Envelope

The envelope was extracted by using a gammatone filter bank of 28 channels with center frequencies between 50 and 5000 Hz. We applied a power law on the absolute values and averaged across the 28 envelopes (same parameters as in Vanthornhout et al. (2018)). These steps were applied because they model the auditory system’s structure (Biesmans et al., 2017).

##### Envelope onsets

The envelope onsets were calculated as the half-wave rectification of the first derivative of the envelope.

##### Phoneme and word onsets

Phoneme and word onsets were coded as dummy variables with a pulse at the beginning of each phoneme, respectively of each word. In order to get there, an aligner (Duchateau et al., 2009) was used to create alignment files containing the timing of each phoneme, respectively each word for the audio files of the stimulus story.

##### Phoneme surprisal

Phoneme surprisal was computed as the negative logarithm of the phoneme prob-ability in the activated cohort. The activated cohort refers to words activated by initial phonemes, e.g., after hearing the sound /pl/, the activated cohort consists of words such as play, plus and plural. Phoneme surprisal is thus a representation of how surprising a phoneme is given the activated cohort. The first phoneme of each word included all words in the active cohort. Phoneme surprisal was calculated based on the SUBTLEX-NL database (Keuleers et al., 2010) and a custom pronunciation dictionary.

##### Phoneme entropy

Phoneme entropy is a measure of the degree of competition between the words congruent with the current phonemic input. For instance, after hearing the sounds /pl/, many possible words are present in the activated cohort (n=999), mirrored in a high degree of competition. Yet, after the next phonemes, the number of possible words decreases (e.g., for /plu/, n=162 and for /plur/, n=19), and thus the activated cohort decreases, reflected in a lower degree of competition. The degree of competition was computed as the Shannon entropy of the words in the activated cohort. Again, the first phoneme of each word included all words in the active cohort. Phoneme entropy was also calculated based on the SUBTLEX-NL database (Keuleers et al., 2010) and a custom pronunciation dictionary.

##### Word surprisal

Word surprisal was calculated as the negative logarithm of the conditional probability of a given word based on the 4 previous words. Word surprisal thus represents how surprising a word is, taking into account the 4 previous words. Word surprisal was calculated using the 5-gram model by Verwimp et al. (2019).

##### Word frequency

Word frequency was calculated as the negative logarithm of the unigram probability and represents how frequently words are used. Given that we used the negative logarithm, words with a higher frequency are reflected in lower scores. Word frequency was also calculated using the 5-gram model by Verwimp et al. (2019).

##### Isolating speech processing levels

An issue when analyzing acoustic and linguistic speech representations is their collinearity, e.g., some top-down linguistic cues at the phoneme and word level are also represented in the speech envelope, such as word boundaries and syllable stress, which contain semantic cues. Vice versa, some bottom-up acoustic information is also present in linguistic phoneme and word level representations (Gillis et al., 2022), e.g., due to amplitude rises and falls defining the boundaries between (pre-)lexical units. Neural signals related to acoustic processing can thus also be captured by neural tracking of linguistic speech representations and vice versa.

In this study, we were interested in disentangling – as far as possible – different speech processing mechanisms. This is important should encoding/decoding modeling be used for diagnosing different language profiles of aphasia in the future, e.g., disentangle individuals that have more problems with acoustic, phonological or semantic processing. Therefore, we regressed out the variance explained by the speech representations that we were not interested in. Specifically, we first applied an ordinary least squares (OLS) regression analysis without regularization, using the EEG signal as dependent variable and speech representations as predictors that share variance with the representations of interest (table 2). All the data was used for training and testing, such that all activity related to the collinear speech representations was regressed out. As a second step, we then used the residual EEG signal of that regression as input for the encoding model in order to model the relationship with the speech representations of interest. The constellation of speech representations used as predictors in the encoding models is illustrated in table 2.

**Table 2:** Constellation of speech representations in the 4 encoding models.

| **Regressing out representations not of interest** | **Boosting on residuals (representations of interest)** | **Encoding model/referred to as** |
| --- | --- | --- |
| phoneme onsets<br>word onsets<br>phoneme surprisal<br>phoneme entropy<br>word surprisal<br>word frequency | envelope<br>envelope onsets | acoustic |
| envelope<br>envelope onsets<br>phoneme surprisal<br>phoneme entropy<br>word surprisal<br>word frequency | phoneme onsets | phoneme-level segmentation |
| envelope<br>envelope onsets<br>phoneme surprisal<br>phoneme entropy<br>word surprisal<br>word frequency | word onsets | word-level segmentation |
| envelope<br>envelope onsets<br>phoneme onsets<br>word onsets* | phoneme surprisal<br>phoneme entropy | phoneme-level linguistic |
| envelope<br>envelope onsets<br>phoneme onsets<br>word onsets* | word surprisal<br>word frequency | word-level linguistic |

\*We decided against regressing out phoneme level linguistic representations for the word level model and vice-versa, because we expected that more data would be needed to get meaningful results from such an analysis.

##### Computation of temporal response function and prediction accuracy

For the encoding analysis of the speech representation of interest, we applied the boosting procedure (David et al., 2007). We made use of the Eelbrain toolbox for this step (https://doi.org/10.5281/zenodo.6992921) (Brodbeck et al., 2021). The data was split into a held-out test set of roughly 2 minutes of the data and a training set containing the rest of the data. The training set was used to perform regression analyses on the residual EEG signal with the speech representation as predictor, for a number of time-shifted versions, resulting in the temporal response function (TRF). The number of time-shifted versions was defined by the chosen integration window length, i.e., –0.1078 to 0.6109 seconds, and the sampling frequency (*(ending time lag-starting time lag)/(1/sampling frequency)*), which resulted in 92 time shifts. The TRF thus provides information about *how* the neural response pattern operates across processing time. This information and the speech representation were then used to predict EEG signals for the test set of the data. The predicted EEG signal was correlated with the originally recorded EEG signal, providing a measure of *how well* the speech representation is encoded in the brain for each electrode, hereafter referred to as prediction accuracy. The higher the prediction accuracy is, the stronger the speech representation is encoded in the EEG signal. This procedure was repeated for different partitions of the data into training and test sets, such that each part of the data was once used as test set, resulting in 12 folds (i.e., k-fold cross-validation). For the TRF, the 12 folds were averaged across to get robust outcome measures. To arrive at the final prediction accuracy, the folds of the predicted EEG signal were concatenated and subsequently correlated with the originally recorded EEG signal.

#### 2.3.4 Determination of the TRF peak latency ranges

To determine in which time windows to perform cluster-based permutation tests, we used the built-in MATLAB function *findpeaks* (MATLAB, 2016) to identify peaks in the TRF. Peak latencies were extracted per speech representation and per participant. An arbitrary range of 150 ms was defined around the average latency of the control group per identified peak (table 3). The average latency per peak of the aphasia group was very similar to the control group (the largest difference among all peaks was 18 ms), such that the selected ranges were valid to find differences between the aphasia and control group. For time ranges that started before 0, the lower bound was set to 0. The 150 ms ranges, as indicated in table 3 and visualized in supplementary figure S.5, were used as time windows to perform cluster-based permutation tests on the TRF (section 2.4 for details). As 26 peaks were found across speech representations, we conducted 26 cluster-based permutation tests.

**Table 3:** TRF peak ranges of the control group used for defining time windows for cluster-based permutation testing.

| Speech representation | Average peak latency (ms) | Range (ms) |
| --- | --- | --- |
| envelope | 29 | 0 - 104 |
|  | 163 | 88 - 238 |
| envelope onsets | 44 | 0 - 119 |
|  | 102 | 27 - 177 |
|  | 209 | 134 - 284 |
| phoneme onsets | 61 | 0 - 136 |
|  | 252 | 177 - 327 |
|  | 305 | 230 - 380 |
| word onsets | 114 | 39 - 189 |
|  | 190 | 115 - 265 |
|  | 308 | 233 - 383 |
|  | 446 | 371 - 521 |
| phoneme surprisal | 221 | 146 - 296 |
|  | 360 | 285 - 435 |
|  | 354 | 279 - 429 |
| phoneme entropy | 243 | 168 - 318 |
|  | 389 | 314 - 464 |
|  | 416 | 341 - 491 |
| word surprisal | 122 | 47 - 197 |
|  | 199 | 124 - 274 |
|  | 320 | 245 - 395 |
|  | 428 | 353 - 503 |
| word frequency | 122 | 47 - 197 |
|  | 197 | 122 - 272 |
|  | 330 | 255 - 405 |
|  | 459 | 384 - 534 |

### 2.4 Statistical analysis

Statistical analyses were performed in R (R Core Team, 2017) and in the Python (Van Rossum and Drake Jr, 1995) toolbox Eelbrain (Brodbeck, 2020).

#### 2.4.1 Group comparison of the strength of neural tracking

For the prediction accuracy analysis, we averaged across all 64 electrodes and conducted a linear model in R in order to investigate group differences and to regress out the covariates, i.e., *average prediction accuracy ∼ group + age + hearing + cognition + alertness + fatigue*. We repeated this for each of the 4 encoding models, i.e. the acoustic, phoneme onsets, word onsets, phoneme and word level model. We checked the normality assumptions using the Shapiro-Wilk test, which failed to reject H0 for each model. The homogeneity of variances assumption was not met in any of the 5 models. Nonetheless, we interpreted the linear models, given that the residuals were normally distributed. As each model was based on a different dataset, we did not control for multiple comparisons in this analysis.

As a second analysis of the prediction accuracies, we conducted a cluster-based permutation test to see if certain electrodes drive the difference between groups. To this end, we used the function *testnd.TTestIndependent* from the Eelbrain toolbox (https://doi.org/10.5281/zenodo.6992921). The cluster-based permutation test is a mass-univariate independent samples t-test that relies on bootstrapping (see Gillis et al. (2021) for more details). We defined a maximum p-value of 0.05 as threshold. We report the number of electrodes in the significant cluster, the v-value and the p-value.

#### 2.4.2 Group comparison of the neural response pattern

To investigate the TRF pattern differences between the control group and the aphasia group, we applied cluster-based permutation tests in the arbitrary integration window ranges identified by the peak latency extraction (table 3). The same function and parameters were used as for the prediction accuracy. As there were multiple peaks identified for each of the 8 speech representations, we corrected the p-value for multiple comparisons within speech representation using the false discovery rate (FDR). Given that this is an exploratory study, we report the uncorrected and corrected significance thresholds.

## 3 Results

### 3.1 Response accuracy to the questions asked during the EEG paradigm

Participants listened to 5 story parts of ca. 5 minutes each while EEG data was recorded. After each part, a yes/no and a multiple choice question were asked about the preceding story part. Figure 3 shows the response accuracy per group separately for the yes/no question and the multiple choice question. For both types of questions, a significant group difference was found (Yes/No question: W = 300, p = 0.01; Multiple choice question: W = 180, p *<* 0.0001). These questions were however not validated and therefore, this result should not be interpreted, but instead be seen as descriptive.

**Figure 3:**
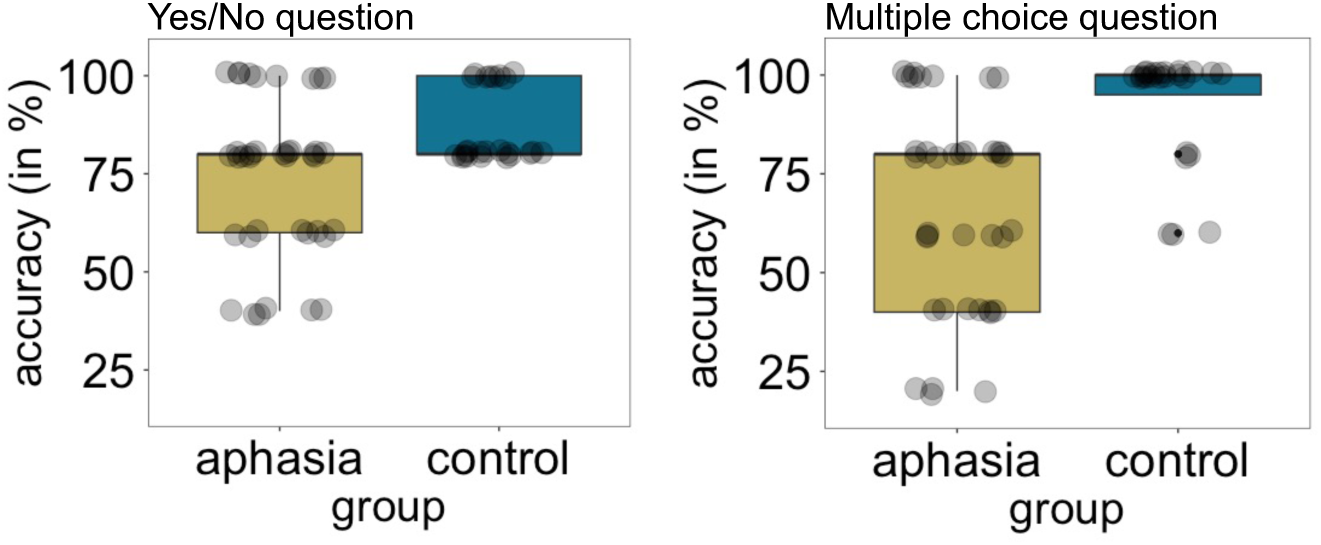
Response accuracy by group for the yes/no questions and the multiple choice questions after each of the 5 story parts that served as stimuli during the EEG measurement. For both types of questions, a significant group difference was found.

### 3.2 Prediction accuracy

When averaging the prediction accuracy across all electrodes, we found a significant group difference in the acoustic model (group effect: F = 7.11, p = 0.01) (fig. 4A). The healthy control group showed higher prediction accuracies in comparison to the aphasia group. The group effect was present despite controlling for the influence of age, hearing, cognition, alertness, fatigue and lesion size in the model. None of the covariates significantly explained the prediction accuracies, except for the acoustic model. Neither the phoneme onset and word onset models nor the phoneme– and word-level linguistic models showed a significant group difference. The results of all models are reported in table 5. Additionally, we also report the results when the covariates are not included in the model (table 4).

**Figure 4:**
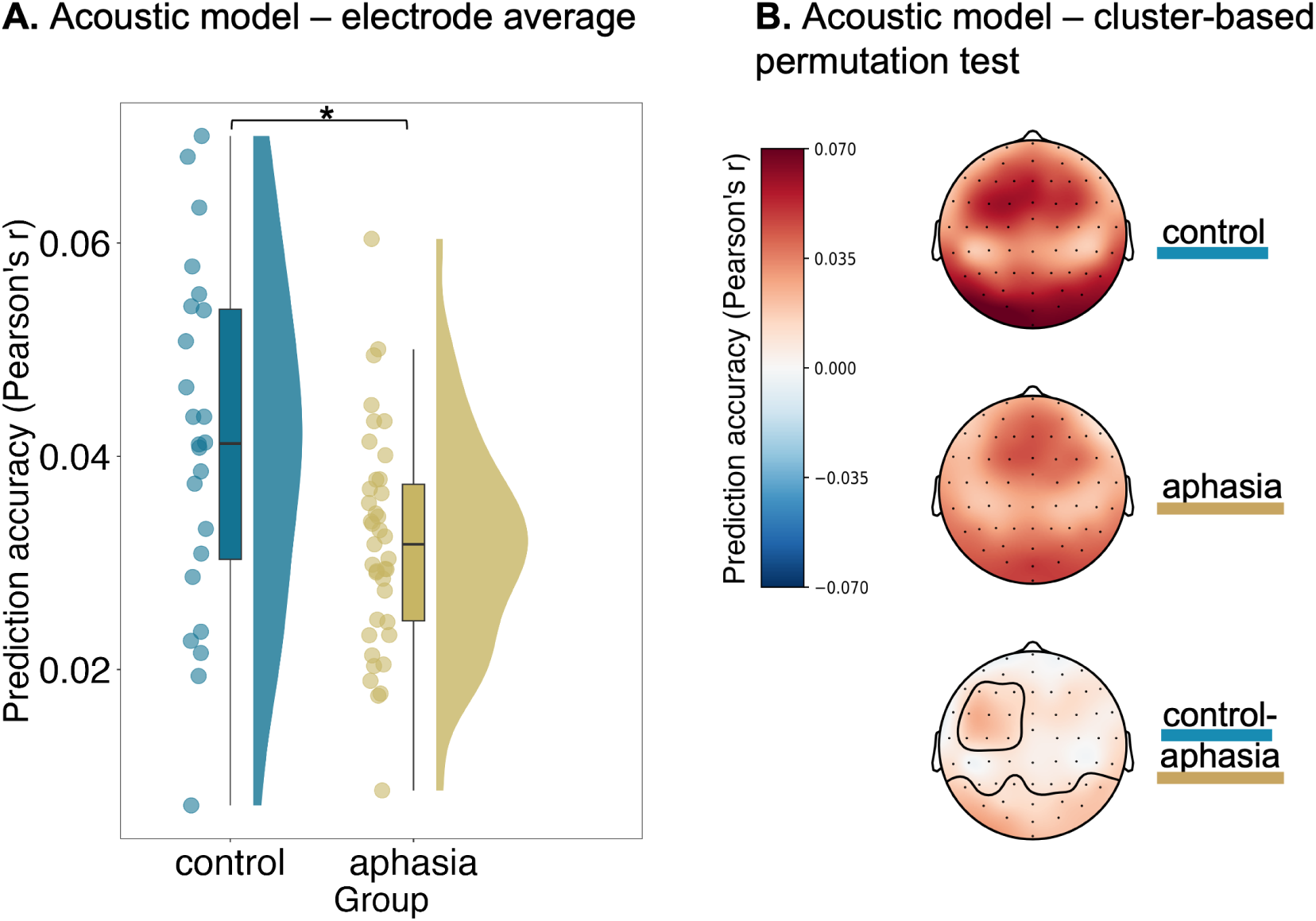
Individuals with aphasia display decreased neural tracking of acoustic speech representations – across all electrodes and in local clusters. **A**. When averaging the prediction accuracies of the acoustic model across all 64 electrodes, we found a significant group difference, even when we controlled for age, hearing, cognition, alertness and fatigue. The acoustic model consists of the speech envelope and its onsets as speech representations. **B.** The cluster-based permutation test revealed 2 clusters that significantly differed between groups, showing lower prediction accuracies in individuals with aphasia. The lowest topoplot consists of the difference between the control group and the aphasia group and the significant clusters are contoured.

**Table 4:** Group comparison results on the prediction accuracy models (without covariates).

| Effect | adj.R<sup>2</sup> | Df | F | Estimate | Standard error | P |
| --- | --- | --- | --- | --- | --- | --- |
| **Acoustic** |  |  |  |  |  |  |
| **Group** | 0.103 | (1,61) | 8.12 | -0.00 | 0.002 | **0.006** |
| **Phoneme onsets** |  |  |  |  |  |  |
| Group | 0.032 | (1,61) | 3.10 | -0.00 | 0.00 | 0.083 |
| **Word onsets** |  |  |  |  |  |  |
| Group | 0.005 | (1,61) | 1.34 | -0.00 | 0.00 | 0.251 |
| **Phoneme-level linguistics** |  |  |  |  |  |  |
| Group | 0.014 | (1,61) | 1.94 | -0.00 | 0.00 | 0.168 |
| **Word-level linguistics** |  |  |  |  |  |  |
| Group | 0.02 | (1,61) | 2.27 | -0.00 | 0.00 | 0.137 |

**Table 5:** Results of the group effect and covariates on the prediction accuracy models.

| Effect | adj.R<sup>2</sup> | DF | F | Estimate | Standard error | p |
| --- | --- | --- | --- | --- | --- | --- |
| **Acoustic** |  |  |  |  |  |  |
| Model performance | 0.083 | (6, 52) | 1.885 |  | 0.013 | 0.1 |
| **Group** |  | 1 | 7.11 | -0.00 | 0.00 | **0.01** |
| Age |  | 1 | 0.16 | 0.00 | 0.00 | 0.69 |
| Hearing |  | 1 | 0.05 | 0.00 | 0.00 | 0.82 |
| Cognition |  | 1 | 0.16 | 0.00 | 0.00 | 0.69 |
| Alertness |  | 1 | 3.58 | 0.00 | 0.00 | 0.06 |
| Fatigue |  | 1 | 0.25 | 0.00 | 0.00 | 0.61 |
| **Phoneme onsets** |  |  |  |  |  |  |
| Model performance | 0.047 | (6, 52) | 1.483 |  | 0.004 | 0.2 |
| Group |  | 1 | 3.06 | -0.00 | 0.00 | 0.08 |
| Age |  | 1 | 0.69 | -0.00 | 0.00 | 0.40 |
| Hearing |  | 1 | 3.09 | 0.00 | 0.00 | 0.08 |
| Cognition |  | 1 | 0.69 | 0.00 | 0.00 | 0.40 |
| Alertness |  | 1 | 0.12 | 0.00 | 0.00 | 0.73 |
| Fatigue |  | 1 | 1.25 | -0.00 | 0.00 | 0.26 |
| **Word onsets** |  |  |  |  |  |  |
| Model performance | -0.038 | (6, 52) | 0.638 |  | 0.004 | 0.69 |
| Group |  | 1 | 1.58 | -0.00 | 0.00 | 0.21 |
| Age |  | 1 | 0.28 | -0.00 | 0.00 | 0.59 |
| Hearing |  | 1 | 0.94 | 0.00 | 0.00 | 0.33 |
| Cognition |  | 1 | 0.26 | -0.00 | 0.00 | 0.61 |
| Alertness |  | 1 | 0.69 | 0.00 | 0.00 | 0.41 |
| Fatigue |  | 1 | 0.08 | -0.00 | 0.00 | 0.77 |
| **Phoneme-level linguistics** |  |  |  |  |  |  |
| Model performance | 0.002 | (6, 52) | 1.024 |  | 0.005 | 0.42 |
| Group |  | 1 | 2.60 | -0.00 | 0.00 | 0.11 |
| Age |  | 1 | 0.00 | -0.00 | 0.00 | 0.97 |
| Hearing |  | 1 | 1.79 | 0.00 | 0.00 | 0.18 |
| Cognition |  | 1 | 0.53 | -0.00 | 0.00 | 0.47 |
| Alertness |  | 1 | 0.36 | 0.00 | 0.00 | 0.55 |
| Fatigue |  | 1 | 0.86 | -0.00 | 0.00 | 0.3568 |
| **Word-level linguistics** |  |  |  |  |  |  |
| Model performance | -0.01 | (6, 52) | 0.903 |  | 0.004 | 0.49 |
| Group |  | 1 | 2.46 | -0.00 | 0.00 | 0.12 |
| Age |  | 1 | 0.46 | 0.00 | 0.00 | 0.49 |
| Hearing |  | 1 | 1.40 | 0.00 | 0.00 | 0.24 |
| Cognition |  | 1 | 0.15 | -0.00 | 0.00 | 0.70 |
| Alertness |  | 1 | 0.26 | 0.00 | 0.00 | 0.61 |
| Fatigue |  | 1 | 0.69 | -0.00 | 0.00 | 0.40 |

DF=degrees of freedom; significant effects are marked in bold.

In order to get an idea how the lesion size affects the link between neural tracking measures and group, we created a hypothetical, statistical scenario in which IWA would have a lesion size of 0. We did this by comparing the intercept of the linear model *average prediction accuracy* ~ *lesion size* including IWA only, to the intercept of the model *average prediction accuracy* ~*group* including both IWA and controls. We found that the 2 intercepts are not significantly different for any of the speech representation models (acoustic: p=0.71; phoneme onsets: p=0.51; word onsets: p=0.80; phoneme linguistic: p=0.72; word linguistic: p=0.64). This means that, if IWA had no lesion (a lesion size of 0), then they would not differ from healthy controls on measures of neural tracking.

The slope coefficient of the model *average prediction accuracy* ~*lesion size* including IWA only was not significantly different from 0 for the prediction accuracy models acoustics, word onsets, phoneme-level linguistics and word-level linguistics (acoustic: p=0.08; word onsets: p=0.59; phoneme linguistic: p=0.13; word linguistic: p=0.59). The phoneme onset model however showed a slope coefficient of lesion size that was significantly different from 0 (phoneme onsets: estimate=-2.35e-05, p=0.01), meaning that a larger lesion size is associated with lower neural tracking scores within the aphasia group, for the phoneme onset model. A relation between lesion size and the phoneme onsets model was also found in a correlation analysis within the aphasia group (phoneme onsets model: Spearman’s r = –0.355, p = 0.026; supplementary fig. S.6).

The cluster-based permutation tests revealed a group difference for the acoustic model, and additionally provided information about what cluster of electrodes is driving the difference between the control group and the aphasia group. Specifically, 2 clusters were found to differ between groups (fig. 4 B). A frontal left-lateralized cluster (number of sensors = 9, v = 25.012, p = 0.02) and a posterior cluster (number of sensors = 15, v = 44.516, p = 0.004) both displayed decreased prediction accuracies in IWA. The phoneme onsets model revealed one cluster that differed between groups (number of sensors = 6, v = 15.511, p = 0.03), namely displaying decreased prediction accuracies in IWA. Neither the word onset model nor the phoneme– and word-level linguistic models showed any significant group differences.

Aphasia is a disorder with a heterogeneous phenotype, as was reflected in the large variability of the language test outcomes (see supplementary table S.1). This made us wonder whether the heterogeneity of language levels in the aphasia group may hide subtle effects in group statistics. Therefore, we decided post-hoc to split up the aphasia group into a more mild and a more severe aphasia group. In the supplementary material (S.1.4), we repeated the group comparison analyses after splitting up the aphasia group. The criteria for splitting up the group is described in more detail in the supplementary material (also see supplementary fig. S.7 and S.10). When averaging the prediction accuracy across all electrodes, we found that the group difference for the acoustic model was not present for the comparison of the control and mild aphasia subgroup, but was present for the comparison of the control group and the aphasia subgroup with more severe language difficulties (supplementary fig. S.8 A). No group effects were observed for any of the other 4 models. The cluster-based permutation test of the acoustic model showed 2 almost identical clusters to the ones found in the analyses with 2 groups (fig. 4 B), however only for the comparison between the control group and the aphasia subgroup with more severe language difficulties (supplementary fig. S.8 B).

### 3.3 Temporal response function

For the speech envelope TRF, we found 2 clusters that differed between the control and aphasia group around 180 ms (fig. 5 A; frontal cluster: time(ms) = [95 243], number of sensors = 28, v = 561.96, p= 0.001; posterior cluster: time(ms) = [142 243], number of sensors = 18, v = –425.69, p = 0.005). The 2 clusters are similar to the clusters from the prediction accuracy of the acoustic model (fig. 4 B). After correction for multiple comparisons via FDR (for n=2 comparisons due to the tested time ranges per speech representations, see table 3), these results remained significant (frontal left: p = 0.002; posterior: p = 0.01). Looking at the neural response to phoneme onsets, we found a cluster that differed between the control group and the aphasia group over frontal electrodes around 280 ms (time(ms) = [235 384], number of sensors = 11, v = 240.58, p = 0.04). This group effect in the neural response to phoneme onsets did not survive correction for multiple comparisons.

**Figure 5:**
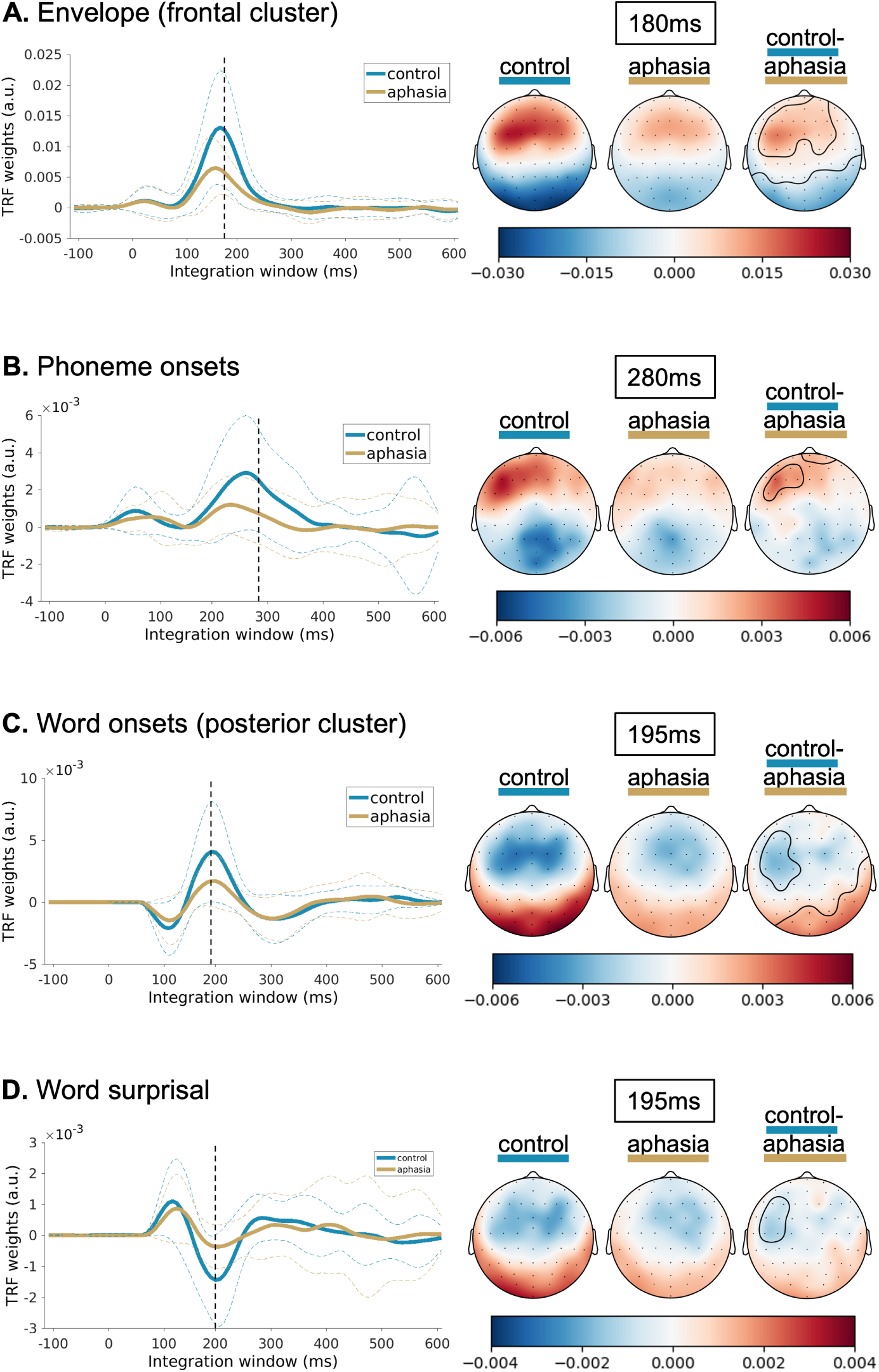
Individuals with aphasia show reduced neural response amplitudes to acoustic and linguistic speech representations in local clusters. In each of the 4 panels, the control and aphasia group average TRFs are plotted. For the left-sided figures, we averaged across the electrodes in the identified clusters. For the right-sided figures, the topoplots are shown at the time point indicated by the dashed line in the left-sided figures. The last topoplot on each panel displays the difference between the control group and the aphasia group, with the clusters being contoured. **A.** The neural response pattern to the speech envelope revealed 2 clusters around 180 ms, a frontal one (visualized) and a posterior one (not visualized). **B.** The neural response pattern to the phoneme onsets revealed a frontal cluster around 280 ms. **C.** The neural response pattern to the word onsets revealed 2 clusters around 195 ms, a frontal one (not visualized) and a posterior one (visualized). **D.** The neural response pattern to word surprisal revealed a cluster with 2 peaks, a posterior peak around 135 ms and a frontal peak around 195 ms (electrodes of this latter peak are visualized). Only the effect of the envelope was robust against correction for multiple comparisons.

No group difference clusters were found for the speech representations envelope onsets, phoneme surprisal and phoneme entropy. However, for the speech representations at the word level, i.e. word onsets and word surprisal, displayed clusters that differed between groups were observed (fig. 5 C and D). The neural response patterns to word onsets displayed 2 clusters that differed between the control and aphasia group around 195 ms (frontal left-lateralized cluster: time(ms) = [134 243], number of sensors = 10, v = –220.3, p = 0.03; posterior cluster: time(ms) = [118 267], number of sensors = 14, v = 267.25, p = 0.016). The neural response patterns to word surprisal displayed a cluster that differed between the control and aphasia group that spread across 2 negative peaks, around 135 ms (difference in posterior left electrodes) and around 195 ms (difference in frontal left electrodes) (time(ms) = [56 204], number of sensors = 21, v = –227.94, p = 0.01). None of these effects survived the correction for multiple comparisons via FDR.

Same as for the prediction accuracy, we split up the aphasia group into a mild aphasia subgroup and an aphasia subgroup with more severe language difficulties and repeated these analyses for exploratory purposes in the supplementary material (supplementary fig. S.9). For the envelope TRF, we found that the control group and aphasia subgroup with more severe language impairments, but not the control and mild aphasia subgroup, showed a significantly different neural response pattern in 2 similar clusters as were found in the group comparison with the full aphasia group (supplementary fig. S.9). This difference in clusters occurred as well around 180 ms, same as in the comparison of the control and full aphasia group. For the envelope onsets TRF, a group difference was found around 200 ms between the control group and the more severe aphasia subgroup. Moreover, the word-level representations also showed significantly different clusters between the control group and more severe aphasia subgroup, but not between the control and mild aphasia subgroup. However, these differences in clusters were found in a later time window than the clusters found in the group comparison between the control and the full aphasia group (200 ms), namely around 360-370 ms. In all 3 speech representations, i.e., word onsets, word surprisal and word frequency, these later clusters occurred over left-lateralized temporal electrodes (supplementary fig. S.9). None of the other speech representations, i.e., phoneme onsets, phoneme surprisal and phoneme entropy, showed any significantly different clusters, in neither of the 2 pair-wise group comparisons.

## 4 Discussion

In this study, we investigated whether IWA display different neural tracking of continuous speech than age-matched healthy controls at multiple processing levels (acoustic to linguistic). To this end, we collected EEG data of 39 IWA and 24 healthy controls while they listened to a continuous narrative. Speech representations were derived from the narrative and their relation to the EEG signal studied. When the neural signals align with the speech properties, this is referred to as neural tracking. This approach allows to explore *how* spatio-temporal neural response patterns operate during continuous speech at multiple processing levels (i.e., TRFs). Further, this method provides a measure of *how well* speech representations are encoded in the EEG data (i.e., prediction accuracy). Concerning prediction accuracies, group differences between IWA and healthy controls were found for processing acoustic speech representations, both located in posterior and frontal clusters (fig. 4). Regarding TRFs, group differences in processing acoustic, segmentation and word-level speech representations were observed at different peaks between 135-280 ms, located in frontal, left-lateralized or posterior regions (fig. 5). However, as there were multiple TRF peaks identified per speech representation, we corrected for multiple comparisons and found that only the clusters identified in the envelope TRF were robust group differences. Nonetheless, as this is an exploratory study, we will discuss all findings here, because they can help to elicit more concrete hypotheses in future investigations.

To date, neural tracking of acoustic and linguistic speech representations has not yet been investigated in post-stroke aphasia. A study in individuals with PPA, a form of dementia that surfaces as language impairment in its initial stages, has however reported increased speech envelope tracking in the theta frequency band compared to healthy controls (Dial et al., 2021). The authors hypothesized that the increased envelope tracking is related to the underlying physiological changes in PPA, i.e., a hypersynchrony between frontal and temporo-parietal cortex as well as a hyperactivity in the frontal cortex (Dial et al., 2021). Given the fundamentally different etiologies of aphasia after stroke and PPA, we did not base any hypotheses on this study. Indeed, we found a rather contrasting pattern of results in the current study, namely decreased acoustic neural tracking in IWA after stroke (fig. 4 A and B). We controlled for the variance explained by segmentation and linguistic speech representations in the acoustic model in order to yield a purer measure of acoustic processing. The observed group difference was also present despite controlling for the variance explained by age, hearing, cognition, alertness and fatigue. Covariates did not significantly explain any part of the variance in the acoustic model.

The envelope TRF confirms the finding of the acoustic prediction accuracy model, namely decreased amplitudes in IWA over left frontal and posterior electrodes, additionally revealing that this difference occurred around 180 ms neural processing time. These results may indicate that IWA track the slow amplitude modulations of speech, which are essential for speech understanding (Shannon et al., 1995; Zeng et al., 2005; Xu and Pfingst, 2008; Oganian and Chang, 2019), to a lesser extent than age-matched healthy controls. Due to controlling the variance explained by the segmentation and linguistic speech representations, the envelope TRF mainly contains the response to acoustic properties of speech. This would mean that IWA also show impaired processing of speech acoustics. This is in line with findings from Kries et al. (2023), who found that 76% of IWA had a low-level acoustic or phonemic processing impairment.

At the sub-lexical and lexical segmentation level, we did not find a significant group difference. However, the TRF analysis revealed locally decreased amplitudes in the neural response to phoneme onsets in IWA at a peak around 280 ms as well as in the neural response to word onsets at a peak around 195 ms. These findings may indicate that IWA have a decreased performance when it comes to parsing continuous speech. Humans segment continuous speech by making use of strategies such as analysis of prosodic contours (bottom-up processes), making use of knowledge about distributional processing of phonological information and about statistical regularities in word-to-object co-occurrences (top-down processes) (Smith and Yu, 2008; Thiessen and Erickson, 2013; Suanda et al., 2014; Fló et al., 2019). Thus, sub-lexical and lexical segmentation of continuous speech is a product of the interplay between bottom-up and top-down processes (David et al., 2007; Shuai et al., 2014; Gaspers et al., 2017). Based on the current findings, these processes that are necessary for speech segmentation may be impaired in IWA.

EEG experiments that examined ERPs have found the N400 to be related to semantic activation and integration into the sentence context (Kuperberg and Jaeger, 2016). N400 studies in IWA reported that IWA in comparison to healthy controls have increased latencies of the N400 (Chang et al., 2016; Kawohl et al., 2010; Khachatryan et al., 2017; Sheppard et al., 2017; Lice and Palmović, 2017) and attenuated amplitudes (Robson et al., 2017; Sheppard et al., 2017; Kielar et al., 2012; Räling et al., 2016; Lice and Palmović, 2017). Given the similarity between the N400 effects and the neural response pattern to word surprisal (Michaelov and Bergen, 2020; Lopopolo and Rabovsky, 2022), we hypothesized to see differences at the linguistic word level (i.e., word surprisal and word frequency) at a time window between 350 and 450 ms. However, we did not find a group difference in the time window of the N400 in the word surprisal and word frequency TRFs. This may be due to the fact that older adults generally seem to have reduced neural tracking of linguistic speech representations (Gillis and Kries et al., 2023). All participants in the current study, healthy age-matched controls as well as IWA, were on average 70 years old. Older adults may use different strategies to process lexical meaning than those captured by word surprisal and word frequency (Federmeier et al., 2002; Wlotko et al., 2010; Spreng and Turner, 2019; Gillis and Kries et al., 2023). More research is needed to determine an ideal set of speech representations to capture semantic processing in healthy older adults, which can subsequently be translated to aphasia research. Another option that may explain why no group difference was found in the target time window of word-level contextual speech properties (i.e., 350-450 ms) is that the heterogeneity within the aphasia group may have masked potential effects. Our exploratory, supplementary analysis with 2 aphasia subgroups – one with milder or compensated language impairments and one with more severe language impairments – displayed significant differences between the control group and the more severe aphasia group in word surprisal and word frequency TRFs at 360-370 ms, thus falling within the N400 time window.

Surprisingly, we found that IWA have decreased amplitudes in the word surprisal TRF around 200 ms. This peak is in line results from Gillis and Kries et al. (2023), where it was present in older, but not younger adults. This peak may be related to lexical segmentation, since the word onsets TRF also showed a group difference at the same latency and over similar electrodes. While we regressed out the influence of word onsets to analyze the response to word surprisal and word frequency and vice-versa, the pulses in the word surprisal and word frequency speech representations were set at the beginning of the word and thus inherently also relate to a certain extent to lexical segmentation.

The IWA that participated in this study all had a left-hemispheric or bilateral lesion caused by stroke and the data was collected in the chronic phase after stroke (i.e., *≥*6 months), the median time that had passed since stroke onset being 16 months. Stroke can cause changes in cerebral blood flow (Rabiller et al., 2015; Brumm et al., 2010) and can create cavities filled with cerebro-spinal fluid at the lesion site (Zbesko et al., 2018; Piastra et al., 2022). Especially the latter mechanism prevails also into the chronic phase after stroke (Piastra et al., 2022; Salinet et al., 2014; Zbesko et al., 2018). Both aspects can impact the conductivity of the neurophysiological activity that is picked up by the EEG, which can lead to asymmetries in topography or changes in specific frequency bands (Vorwerk et al., 2014; Cohen et al., 2015; Park et al., 2016; Cassidy et al., 2020). Therefore, in the following paragraph, we will discuss whether the decreased neural tracking effects that we observed in IWA may be related to these underlying anatomical changes that occur after stroke and influence the EEG signal or whether they are indeed a trace of decreased processing of speech.

Many of the clusters that differed between healthy controls and IWA occurred over left-sided electrodes (fig. 4 and 5). This region coincides with the region that is impacted by left-hemispheric lesions in the area supplied by the middle cerebral artery, which is the lesion location for 79% of IWA in this study (fig. 1; supplementary table S.1). Thus, the lesion and hence, the altered conductivity, may influence the EEG signal over these electrodes. While the EEG analysis method that we used here is not a direct measure of the raw EEG signal – as is the case for ERPs – the altered conductivity may still affect measures of neural tracking over lesion sites. The prediction accuracies are computed as the correlation between the recorded EEG signal and the EEG signal predicted via modeling of the TRF and the speech representation. This correlation does not take into account the absolute amplitude of the signals it compares. However, since the signal-to-noise ratio of the EEG signal recorded over the lesion site is most likely lower than for other electrodes, the prediction it makes will be more noisy too, resulting in a lower prediction accuracy. TRF amplitudes are additionally more directly influenced by the recorded EEG amplitude. Thus, theoretically neural tracking of speech properties can be impacted by the lesion-induced changes in conductivity of the neurophysiological activity. We tested whether we would find a group difference when the lesion size is regressed out for the aphasia group (section 3.2). We found that if IWA had no lesion, then they would not differ from controls. We also found for the phoneme onsets model that the larger the lesion size is, the lower the neural tracking scores within the aphasia group are. Together with the main results, this indicates that the altered conductivity over the stroke lesion site, and in part probably also a difference in processing slow amplitude modulations of speech may jointly explain the lower neural tracking in the aphasia group. It should however be noted that this study was not designed to determine what factors cause the group difference in neural tracking.

Some of the clusters in which the neural response amplitude significantly differs between groups (fig. 4B and 5) occurred over posterior electrodes. We can only speculate about these posterior clusters. As shown in supplementary section S.1.5, IWA with a posterior lesion do not explain the occurrence of the posterior clusters (supplementary fig. S.11). One possibility is that the large lesion overlap of IWA in the left inferior frontal gyrus (fig. 1), mixed with lower speech processing-related activity in those areas, led to lower measures of neural tracking over those electrodes. This resulted in a slight asymmetry with higher amplitudes over anterior right-sided electrodes. The control group on the other hand showed a slight asymmetry with higher amplitudes over anterior left-sided electrodes. These asymmetries in both groups slightly change the dipole orientation in opposite directions. When the aphasia group topography is subtracted from the control group topography, it consequently results in larger differences over anterior left-sided electrodes and over posterior right-sided electrodes.

## 5 Conclusions and future outlook

In sum, our results show that measurements of neural tracking to specific speech properties may be a promising avenue for future diagnostic and therapeutic applications. Especially the decrease in IWA in processing of acoustic cues, such as the amplitude fluctuations and the onsets of phonemes and words, seems to be a robust effect. Future studies may confirm the potential role of neural tracking of acoustic and linguistic speech representations to provide profiles of language processing difficulties in IWA (i.e., acoustic, phonological, semantic). However, confounding factors, such as effects of the stroke lesion on the EEG signal need to be taken into account in future investigations of neural tracking of speech in aphasia. An interesting approach to address this issue could be the recruitment of a control group consisting of individuals with a stroke without aphasia.

Once neural tracking in aphasia is better understood, the application potential of this method could be manifold. A study that is currently in preparation found that IWA can be distinguished from healthy controls with 83% accuracy based on neural envelope tracking with mutual information of only 5 to 7 minutes of EEG data (De Clercq et al., 2023). Looking a step further into the future, an aphasia diagnosis based on processing levels of different speech representations could complement behavioral diagnosis and inform speech-language pathologists further which functions should be trained in therapy. Test-retest practice effects (i.e., improved performance on repeated tests due to remembering items or training test-specific skills) could be avoided during therapy follow-up. Moreover, the method could be useful in the acute phase after stroke, when behavioral diagnostic tests are too exhausting for patients. This would still have to be tested in a clinically more compatible experimental setting in the future, but work by De Clercq et al. (2023) shows that only few minutes of recording time would be needed to get reliable data. Additionally, articulatory speech representations (e.g., mouth aperture, tongue protrusion (Mitra et al., 2010)) could be investigated to analyze effects of production impairments in aphasia during listening. Further, studying neural processing during speech production, using the same analytical framework, also offers a possibility to study fluency impairments in IWA. Due to the feasibility of EEG in the clinical context, the efficiency of the paradigm and the versatility of applications, examining neural tracking of naturalistic, continuous speech provides a powerful approach to studying aphasia.

## Acknowledgements

The authors would like to thank all participants, especially all the brave participants with aphasia and their partners, family or friends that support them. Furthermore, the authors would like to thank Dr. Klara Schevenels for helping with recruitment of participants with aphasia. Thanks also to Dr. Toivo Glatz for methods advice. Moreover, the authors would like to thank everyone who helped with data collection and recruitment: Janne Segers, Rosanne Partoens, Charlotte Rommel, Dr. Ramtin Mehraram, Ines Robberechts, Laura Van Den Bergh, Anke Heremans, Frauke De Vis, Mouna Vanlommel, Naomi Pollet, Kaat Schroeven, Pia Reynaert and Merel Dillen.

## Data availability statement

All data generated or analyzed for this study are available on Open Science Framework (OSF) under the following link: https://osf.io/6t2ja/.

## Ethics approval and patient consent statement

The study received ethical approval by the medical ethical committee of KU Leuven and UZ Leuven (S60007) and is in accordance with the declaration of Helsinki. Informed consent was obtained from all participants for the recruitment via screening and for the data collection in the chronic phase.

## Author contribution statement

Conceptualization: MV, JK, TF; Investigation: JK, PDC, MV; Project Administration: JK; Data Curation: JK; Resources: MV, TF, RL; Methodology: JK, MV, JV, MG, PDC, TF; Formal analysis: JK, MG, JV, PDC; Visualization: JK, PDC, MG; Writing – Original Draft Preparation: JK; Writing – Review and Editing: JK, PDC, MG, JV, RL, TF, MV; Supervision: MV, TF, RL; Funding Acquisition: MV, JK

## Supplementary material

### S.1.1 Recruitment, data collection and participant details

#### S.1.1.1 Flowchart of recruitment procedure

**Figure S.1:**
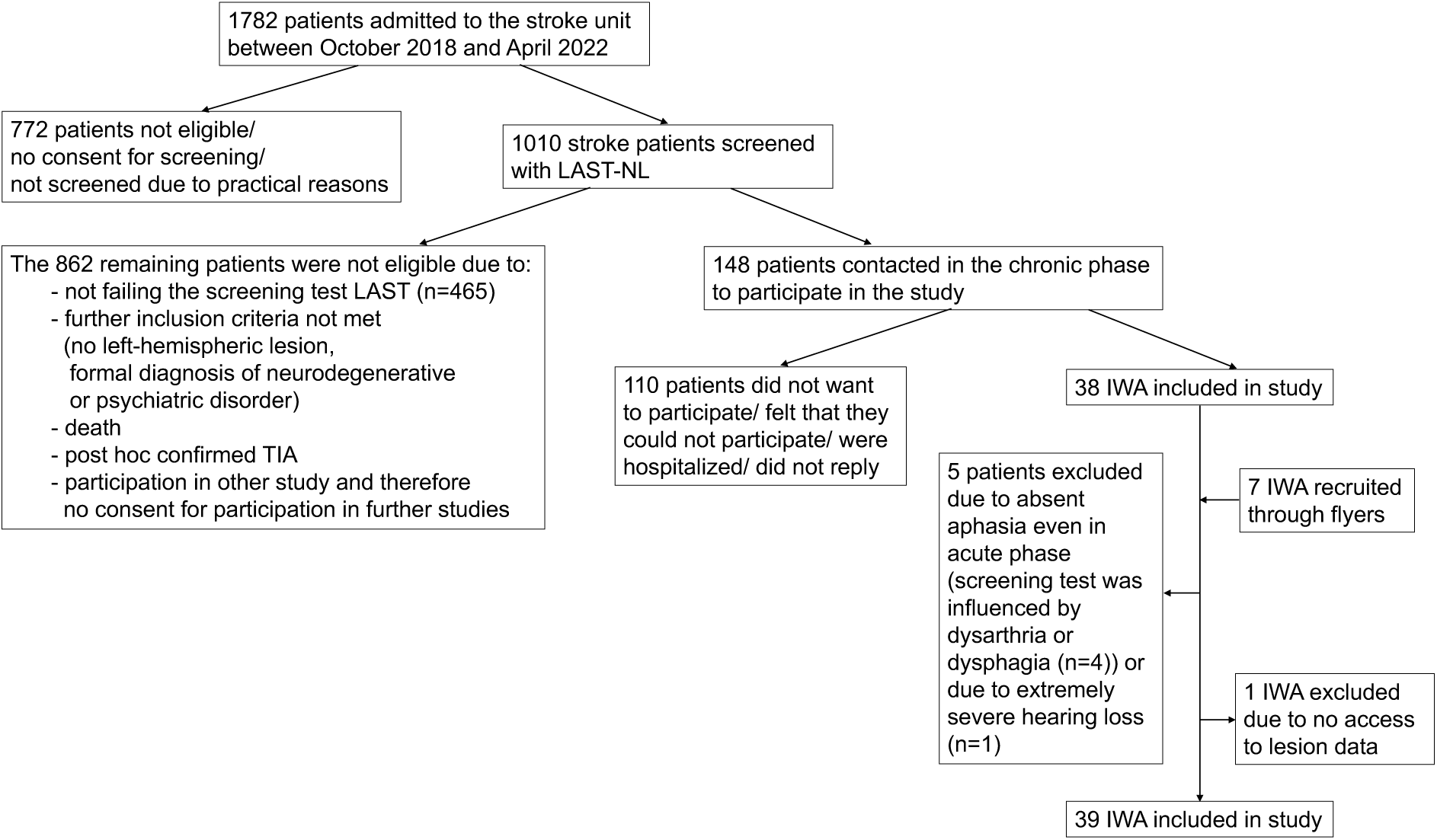
Flowchart of procedure to recruit participants with aphasia.

#### S.1.1.2 Timeline of data collection

Data collection had to be stopped during winters of 2020/2021 and 2021/2022 due to the COVID-19 pandemic and thus took place mainly between the months June and October of each year. Healthy controls and participants with aphasia were recruited in alternation throughout the 3 years, which allowed us to age-match healthy controls to the IWA. Moreover this ensured that changes in EEG hardware did not influence one group more than the other. The electrode set was changed in September 2021 to prevent future problems due to extensive usage of the previous set.

**Figure S.2:**
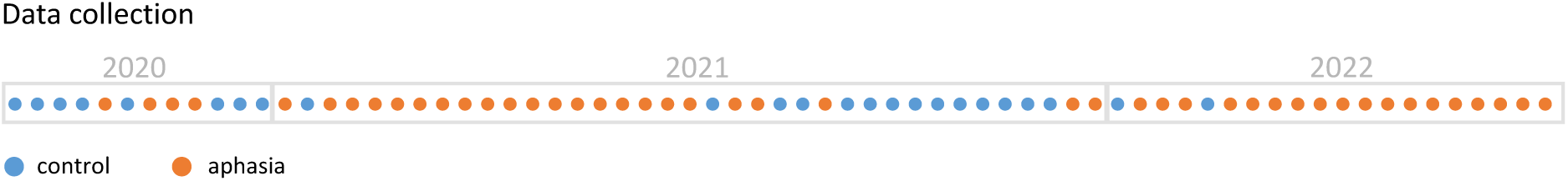
Timeline of data collection of healthy controls and participants with aphasia.

#### S.1.1.3 Participant details

**Table S.1:**
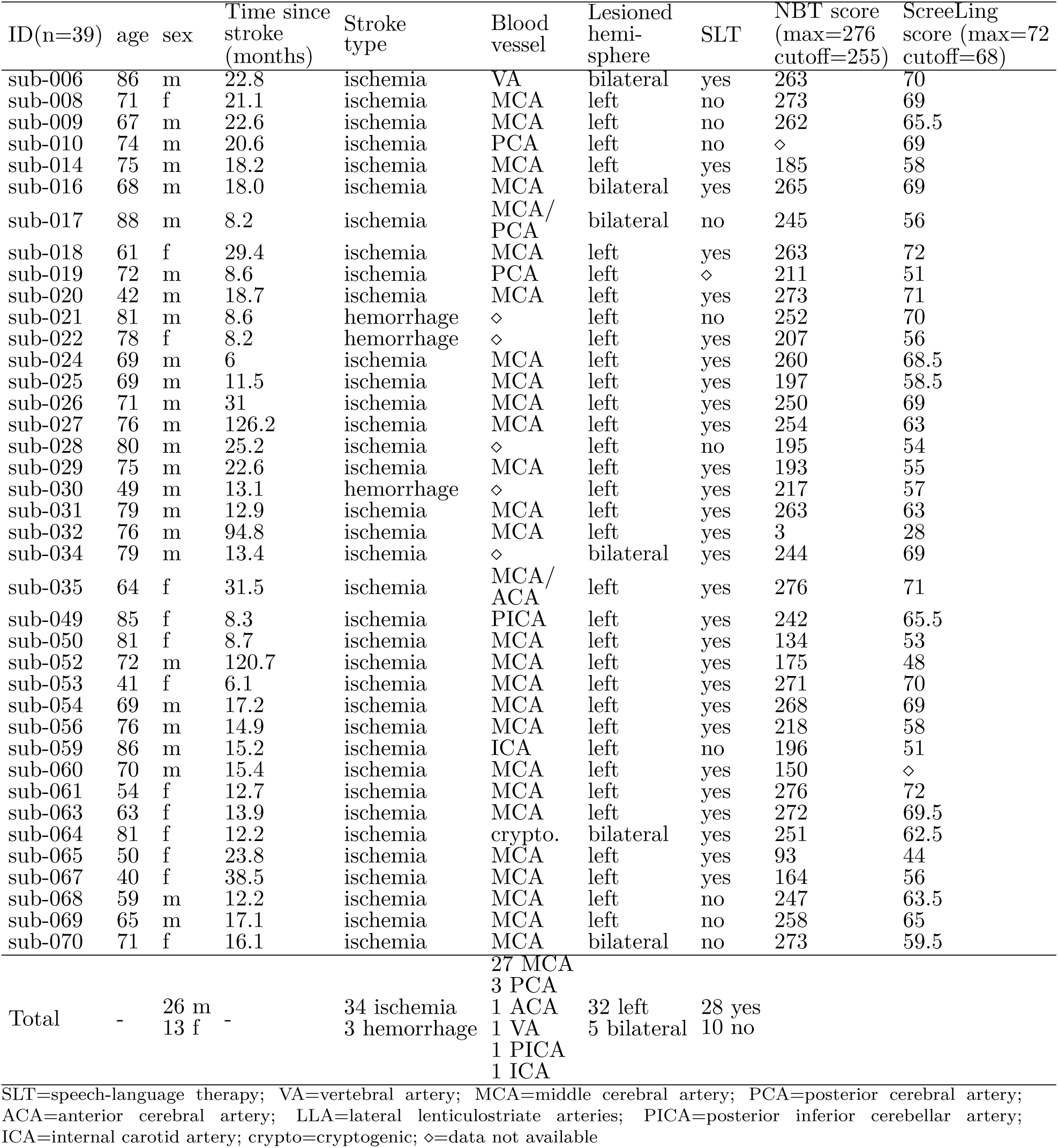
Demographic, lesion and diagnostic information of the aphasia group.

**Figure S.3:**
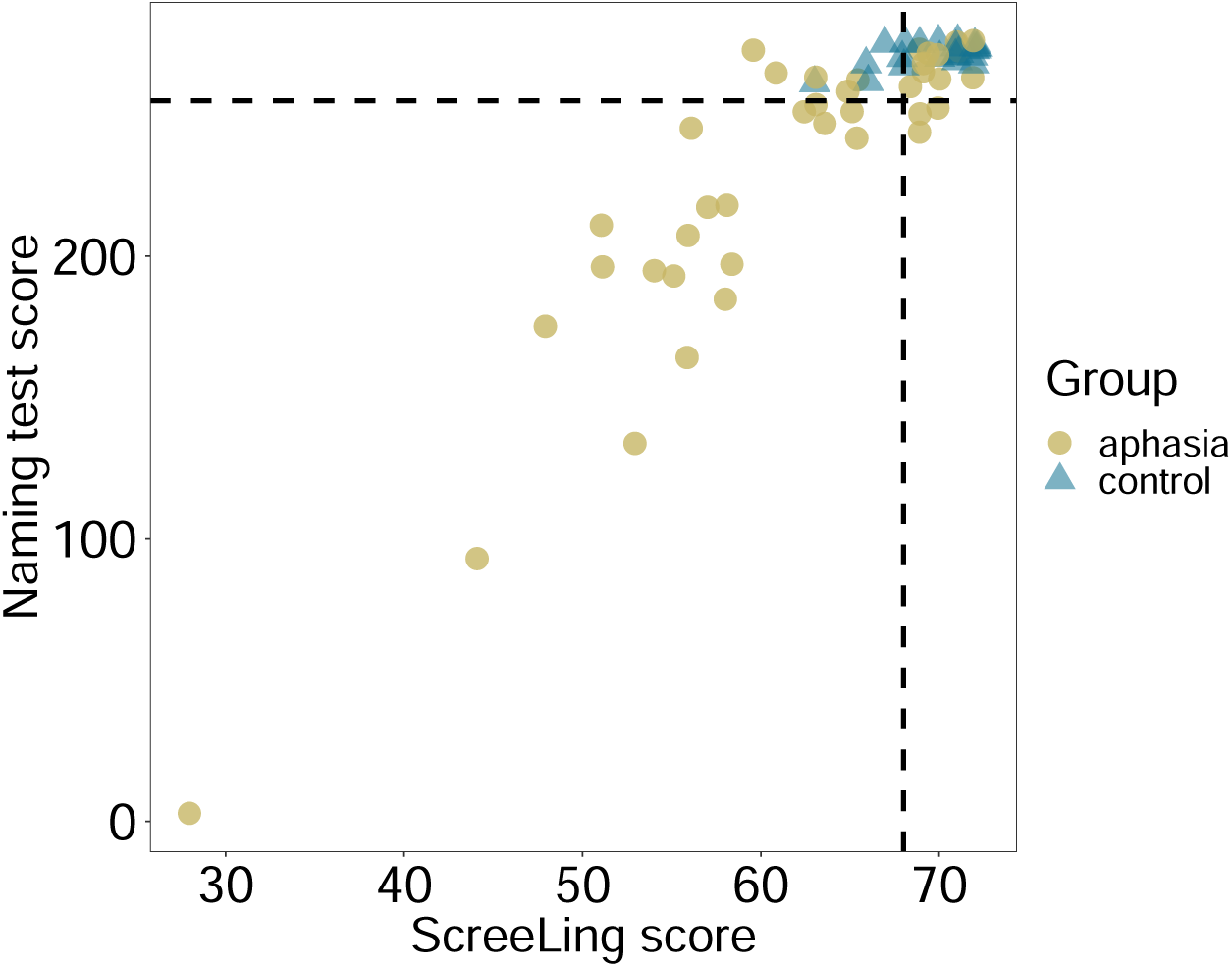
Visualization of scores on language tests. The cut-off scores are marked by the dashed lines.

#### S.1.1.4 Lesion delineation

MRI data collection was not a part of the planned data collection for this study, but ethical agreement was given to access the scans from the medical files of the university hospital. Thus, we had access to T2-weighted FLAIR images that were administered in the acute stage post-stroke, based on which the segmentations of the affected stroke tissue were manually performed. For 10 participants, scans were available from the chronic phase post-stroke, as they took part in an MRI study (unpublished). 12 participants had participated in an earlier aphasia study, and thus we could use the already established segmentation maps.

#### S.1.1.5 Monitoring alertness and fatigue throughout the experimental session

The subjective ratings of alertness and fatigue were administered in order to use them as covariates in statistical models. The EEG measurement took place between t1 and t2, therefore we averaged across t1 and t2 of the alertness and fatigue rating respectively, which were then used as covariate factors for the analysis of prediction accuracy models (fig. 4). We also analyzed the course of alertness and fatigue ratings between groups. Linear mixed effects models were employed for this purpose. The analysis of the alertness question revealed a main effect of group (p*<*0.001), but no main effect of time point or interaction effect (time point: p=0.06; interaction: p=0.226; fig. S.4 C). Thus, across time points, individuals with aphasia were less alert than healthy controls. The analysis of the fatigue question showed significant main effects (group: p*<*0.001; time point: p=0.015) and a significant interaction effect between group and time point (p=0.02; fig. S.4 D). Post hoc testing revealed that while the initial level of fatigue was at a similar level for both groups at the beginning of the test session (t1; p=0.92), individuals with aphasia were more tired than healthy controls after the first part of the experimental session (t2; p*<*0.001) and at the end of the protocol (t3; p=0.002).

We offered IWA to continue the administration of the second part of the experimental protocol on another day in case they felt too exhausted after the EEG measurement (fig. S.4 E). Indeed, 5 out of 41 IWA felt too exhausted at t2 and chose to do the rest of the tests on another day. This second session took place at the patients’ homes, so that they did not have to arrange transportation to the lab again.

**Figure S.4:**
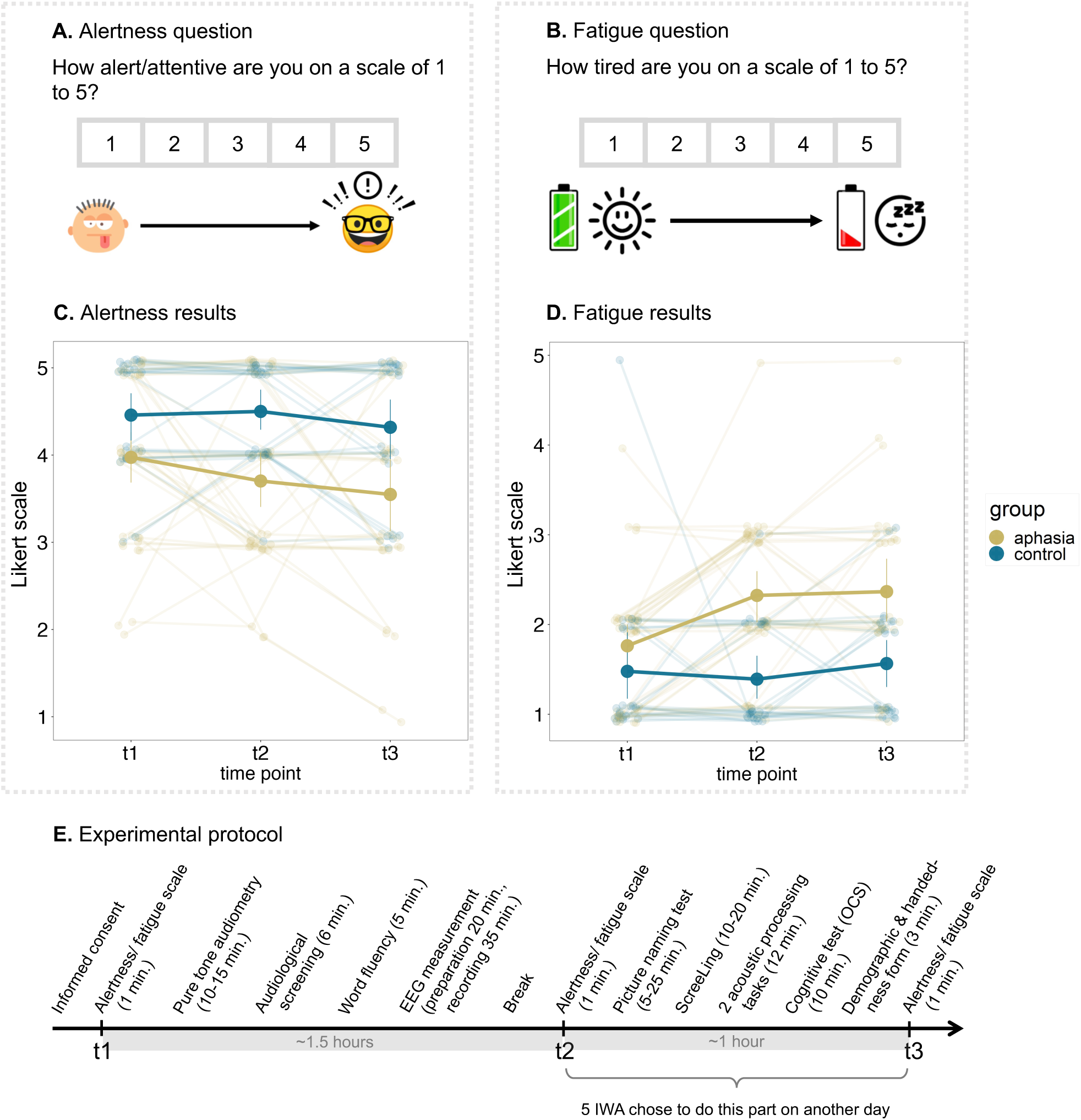
Subjective ratings of alertness and fatigue throughout the experimental session. **A. and B.** The alertness and fatigue questions as presented on a tablet to the participants, with visual icons to aid comprehension of the task. The questions were simultaneously presented auditorily. **C.** A linear mixed effects model showed a main effect of group, but no main effect of time point or interaction effect for the alertness scale. **D.** A linear mixed effects model showed a significant interaction effect between group and time point for the fatigue scale. **E.** Visualization of the experimental protocol with regard to the time points at which the alertness and fatigue scales were administered.

### S.1.2 Determined TRF peak latency ranges

**Figure S.5:**
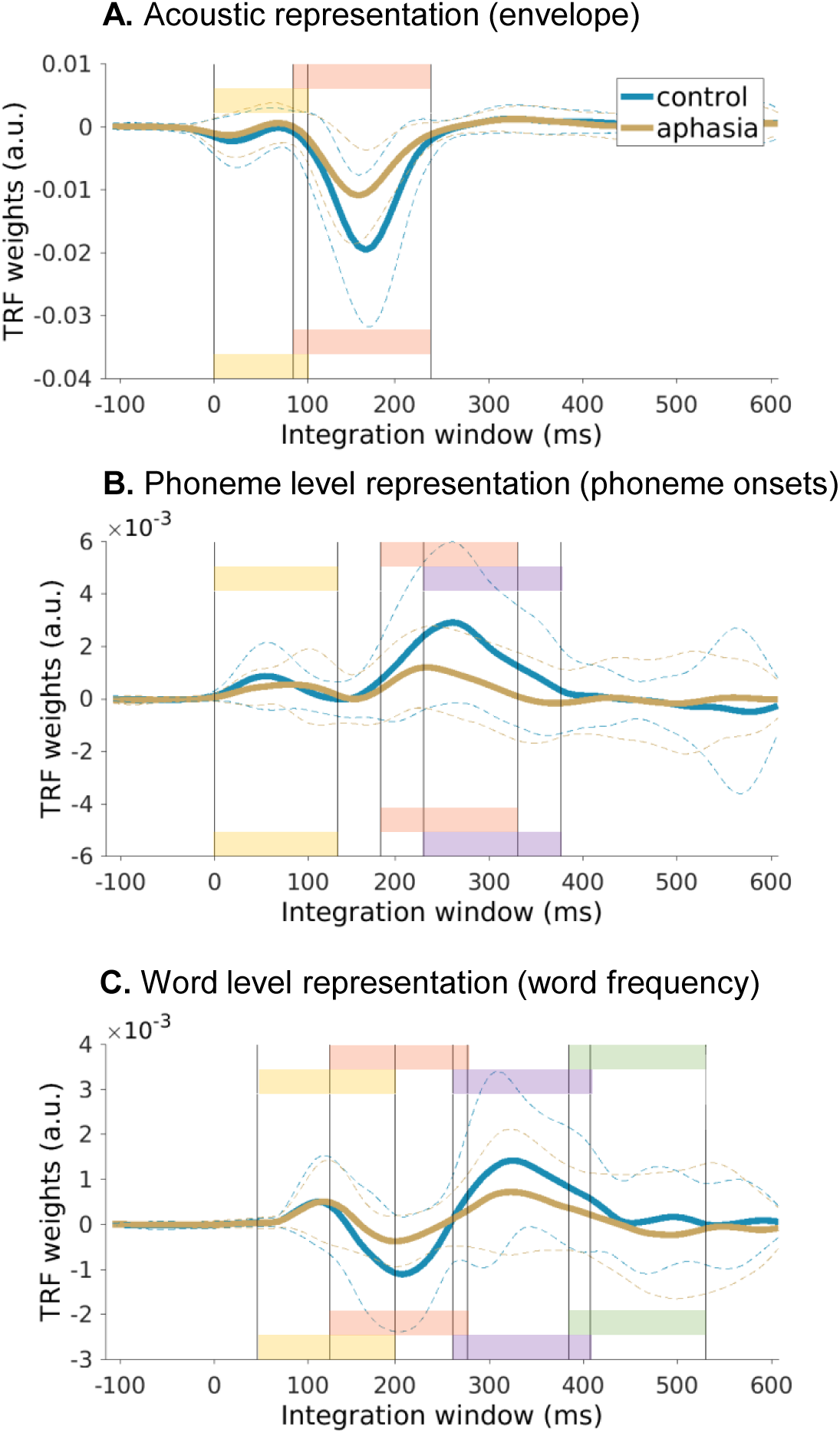
Time ranges of 150 ms visualized in 3 speech representations.

### S.1.3 Within-aphasia group exploratory correlations between prediction accuracy and behavioral measures

**Figure S.6:**
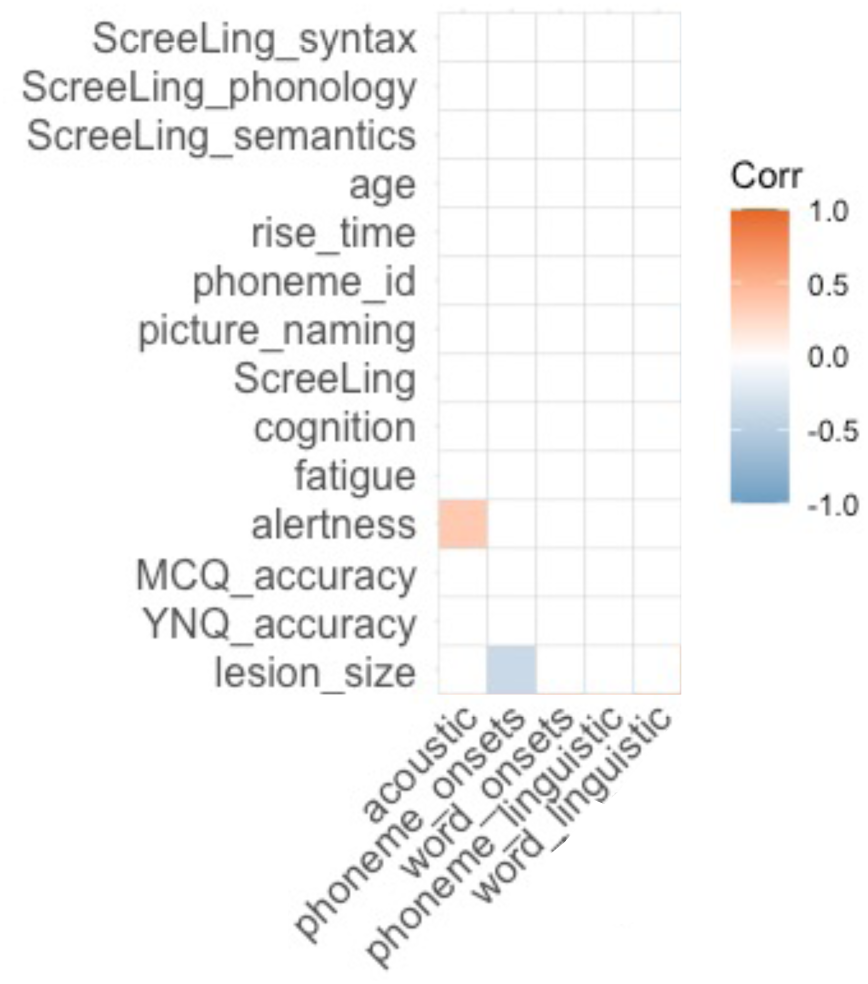
Exploratory correlation matrix. Only significant correlations are colored. Note: Rise time and phoneme identification tasks are measuring acoustic and phoneme processing, the tasks are described in detail in (Kries et al., 2023). MCQ=multiple choice question; YNQ=Yes/No question.

The phoneme onset prediction accuracy model is negatively correlated with lesion size: The higher the prediction accuracy, the smaller the size of the lesion.

The acoustic prediction accuracy model is positively correlated with alertness: the higher the prediction accuracy, the higher the alertness.

### S.1.4 Comparing the control group to a mild aphasia subgroup and an aphasia subgroup with more severe impairment

The aphasia group in the main text consists of 39 IWA who all had language problems at least in the acute phase after stroke, as measured by scoring below the cut-off on a standardized diagnostic test (Comprehensive Aphasia Tes-NL, ScreeLing, Aachen Aphasia Test). Aphasia is a disorder with a heterogeneous phenotype, as was indeed mirrored in the large variability of the language test outcomes (see table S.1). This made us wonder whether subtle effects may be hidden by group statistics due to this large variability. Therefore, we decided post-hoc to split up the aphasia group into a more mild and a more severe aphasia group. We based the splitting on the outcomes on the diagnostic test ScreeLing (max. score: 72; cut-off score: 68) and the picture-naming test NBT (max. score: 276; cut-off score: 255) (fig. S.7), which resulted in a mild aphasia subgroup of 18 participants and an aphasia subgroup of 21 participants. The control group consisted of 24 participants.

Two inclusion criteria for the mild aphasia subgroup were formulated, i.e., not scoring below cut-off on either of the 2 tests at the time of data collection or scoring below cut-off on only one test, as long as the ScreeLing score is within the range of controls (range: 63-72). The reason to set this extra criterium for the ScreeLing, but not for the NBT, is that the ScreeLing is a diagnostic test characterizing language impairment across 12 different tasks that involve phonological, semantic and syntactic processing. Failing on the ScreeLing below the range of control participants is a clear indication of language impairment. The NBT on the other hand only tests one specific symptom of aphasia, i.e., anomia, and does not reflect a holistic assessment of language performance. Nonetheless, we included the NBT for categorizing the aphasia group, because anomia is present to varying extents across aphasia subtypes. Please note, the fact that IWA scored above cut-off threshold on both language tasks does not necessarily mean that they are recovered and do not have aphasia anymore. Many of these IWA were still in therapy at the moment of data collection and might have done the same tests at earlier times during therapy or might have learned compensation strategies in therapy. They had a left-hemispheric lesion after stroke and they had aphasia at least in the acute phase after stroke.

**Figure S.7:**
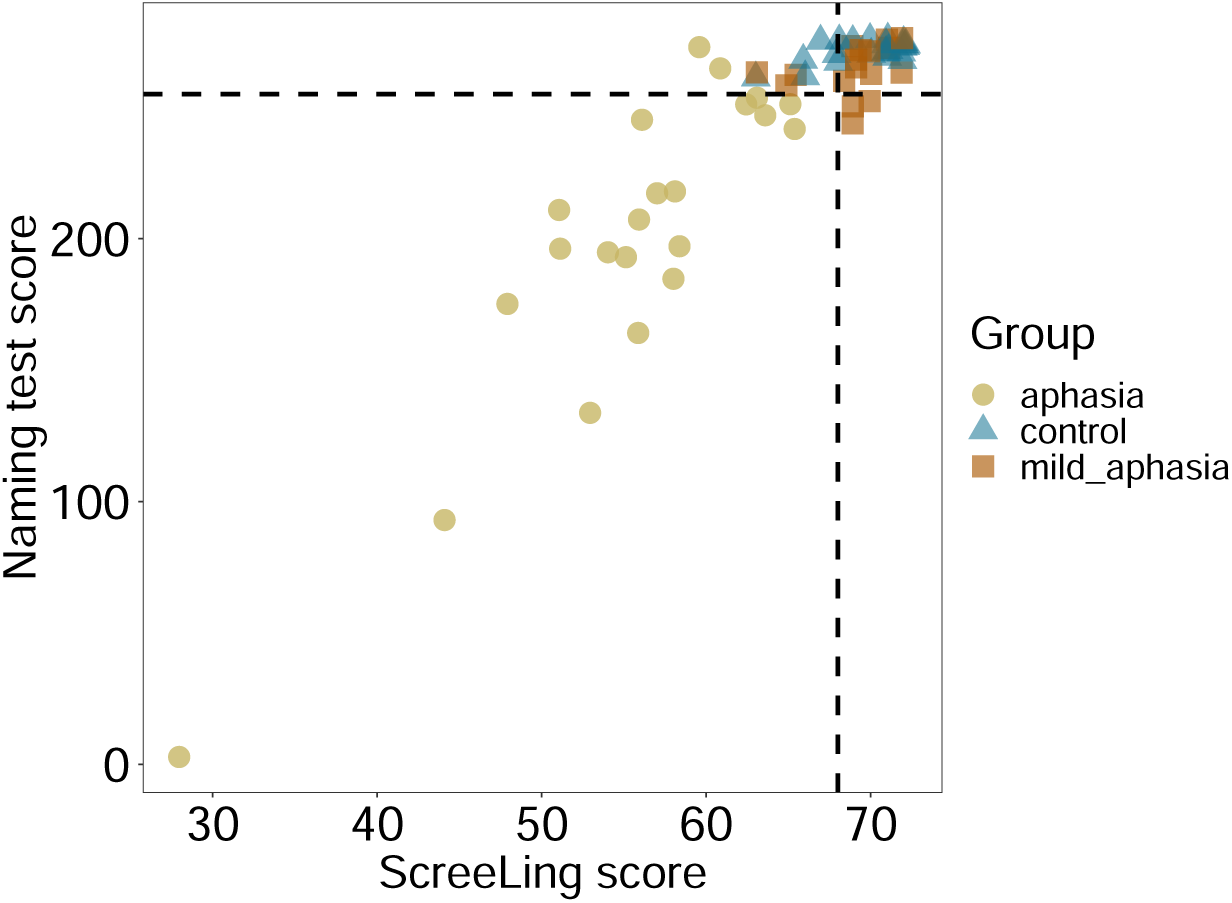
Visualization of scores on language tests used for splitting up the aphasia group into a milder and a more severe group.

We repeated the same analyses for these 3 groups (control, mild aphasia, aphasia) as for the 2 groups in the main text (control, total aphasia group), i.e., group comparisons of the model prediction accuracies averaged over all electrodes, cluster-based permutation tests of the topography of the prediction accuracy and cluster-based permutation tests of the TRFs. These analyses were not corrected for multiple comparisons.

#### S.1.4.1 Prediction accuracy

We found a significant main effect of age for the acoustic model (F(2) = 3.69; p = 0.031) (fig. S.8 A). Post-hoc pairwise comparisons revealed that there is a difference between the control and aphasia subgroup (p = 0.032), but not between the control and mild aphasia subgroup. We controlled for the influence of age, hearing, cognition, alertness and fatigue. None of the other neural tracking models showed any significant group effect.

For the acoustic model, we found 2 significant clusters for the comparison between the control and aphasia subgroup (anterior left-lateralized cluster: v = 32.728, p= 0.014; posterior cluster: v = 37.629, p = 0.009), but none for the comparison between the control and mild aphasia subgroup (fig. S.8 B). The cluster locations are in line with the clusters found for the group comparison between the control group and the full aphasia group (fig. 4). None of the other models revealed any significant clusters between groups.

**Figure S.8:**
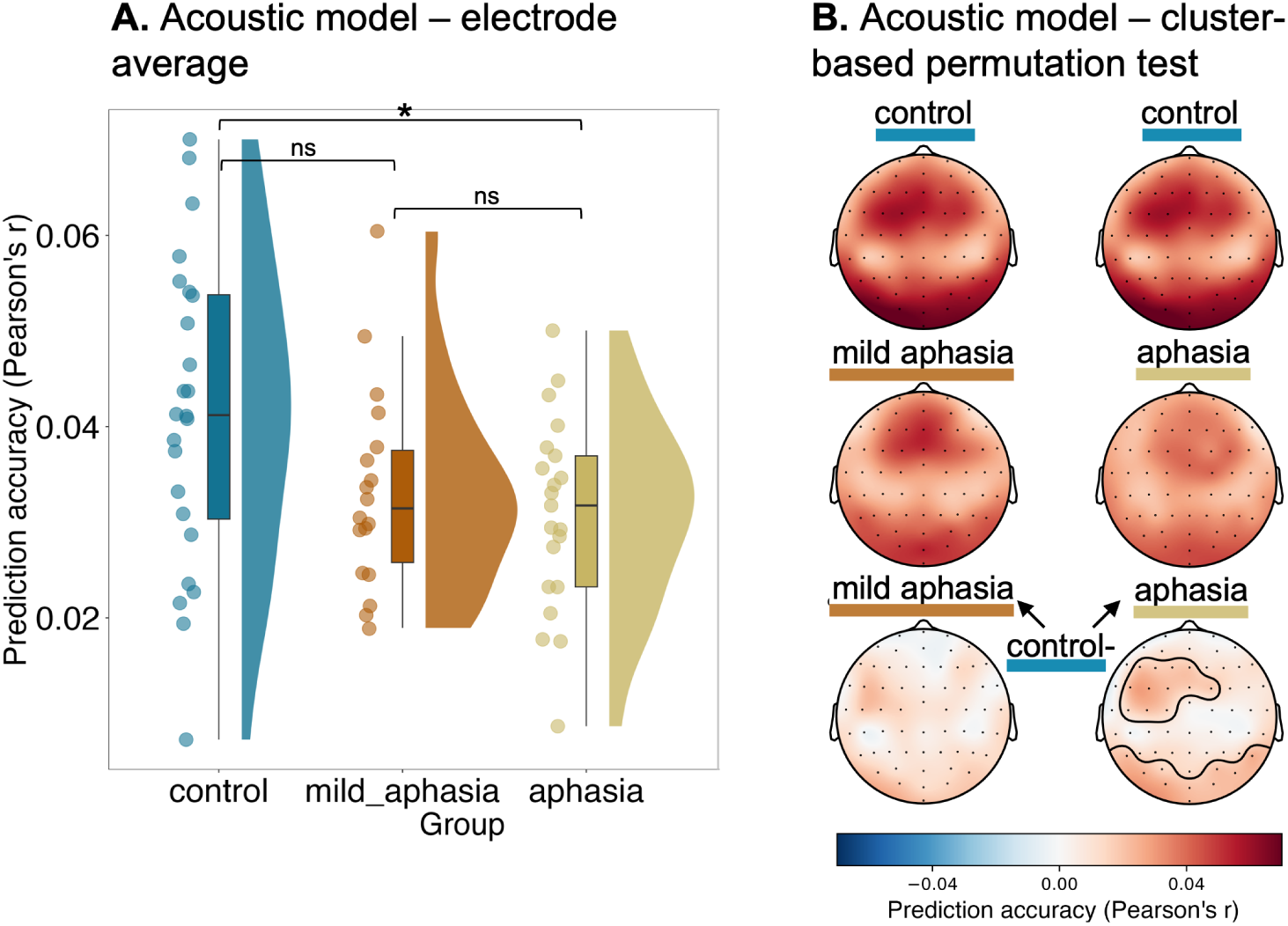
Only the subgroup of aphasia with more severe language difficulties shows decreased neural tracking of acoustic speech representations across all electrodes and in local clusters. **A**. When averaging the prediction accuracies of the acoustic model across all 64 electrodes, we found a significant group effect, even when we controlled for age, hearing, cognition, alertness, fatigue and lesion size. The acoustic model consists of the speech envelope and its onsets as speech representations. **B.** The cluster-based permutation test for the group comparison between control and aphasia subgroup, but not between the control and mild aphasia subgroup, revealed 2 clusters that significantly differed between groups, showing lower predicition accuracies in individuals with more severe aphasia. The lowest topoplot consists of the difference between the control group and the aphasia/mild aphasia group and the contours encircle the significant electrode clusters.

#### S.1.4.2 Temporal response functions

For the speech envelope TRF, we found 2 significant clusters for the comparison between control and aphasia subgroup around 180 milliseconds (ms) (anterior left-lateralized cluster: v = 384.99, p= 0.007; posterior cluster: v = –271.52, p = 0.034), but no significant clusters for the comparison between the control and the mild aphasia subgroup. The 2 significant clusters were located anteriorly left-lateralized and posteriorly, which is in line with the results from the TRF cluster-based permutation test between the control group and the full aphasia group in the main text (fig. 5), but also in line with the results from the prediction accuracy of the acoustic model (fig. 4 and S.8). Note that the time window of this cluster is also in line with the results in the main text (fig. 5).

For word-level segmentation and linguistic speech representations, we also found differences between the control group and the aphasia subgroup, but not between the control group and the mild aphasia subgroup (fig. S.9). However, the time points of the significant clusters that were found are not the same as were found in the main text (5). In the main text (comparing the control group and the full aphasia group), we found clusters in the neural response around 200 ms for word onsets, word surprisal and word frequency, whereas for the comparison between the control group and the aphasia group with more severe difficulties, we found clusters in the neural response around 360 ms for the same speech representations (fig. S.9). For the neural response to all 3 representations, the clusters that differ between groups around 360 ms are located over temporal left electrodes.

For the other speech representations, no significant clusters were found.

**Figure S.9:**
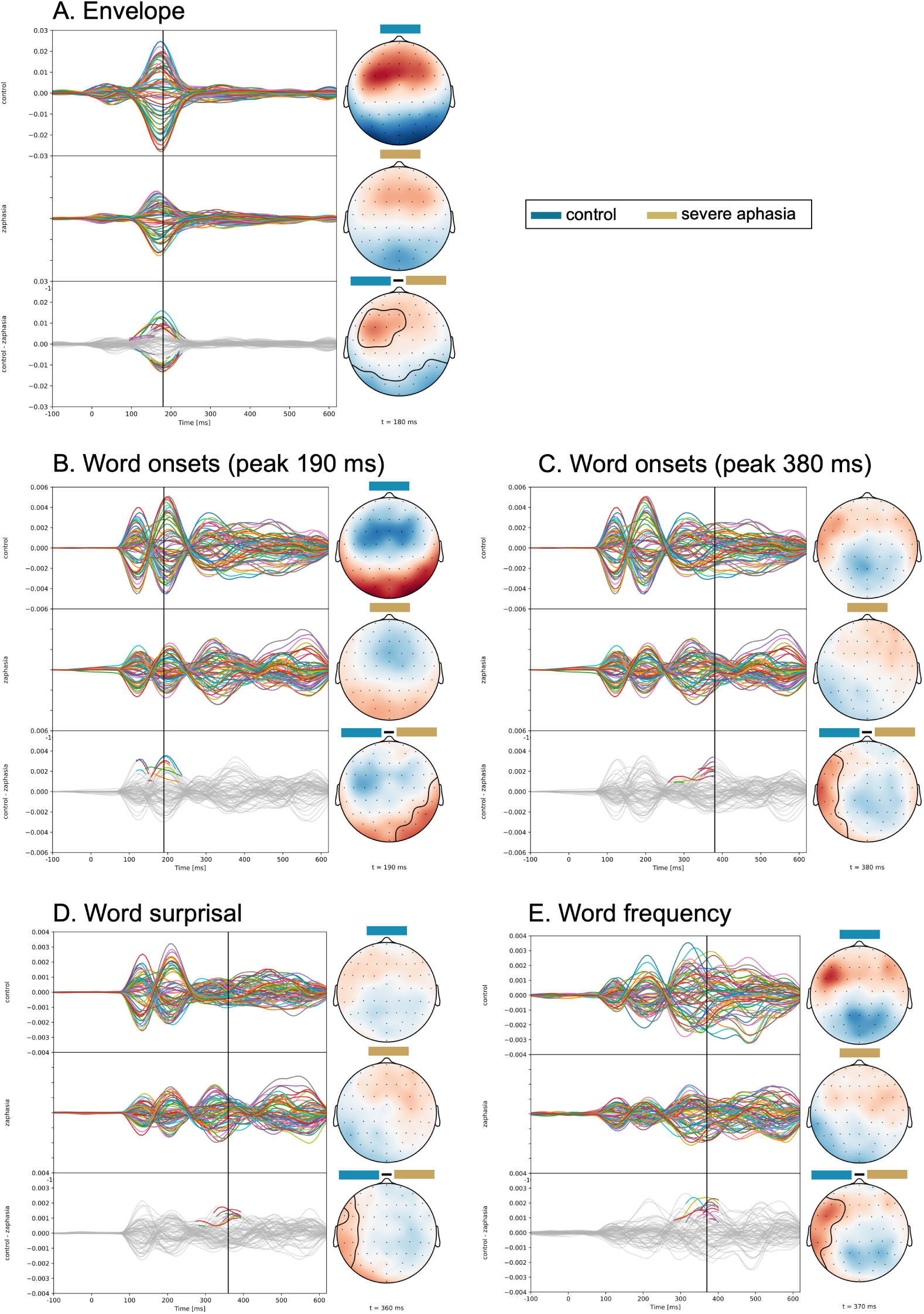
Individuals with more severe aphasia show reduced neural response patterns to acoustic and linguistic representations in local clusters compared to controls. The top most topoplot of each panel shows the control group average and the middle plot shows the aphasia subgroup average. The spaghetti plots show the TRF weights across the integration window. Each line represents one of the 64 electrodes. The topoplots display the topography of the neural response at the time point indicated in the TRF plot with a vertical black line. The bottom most TRF plot and topoplot display the difference between the control group and the aphasia subgroup with more severe language problems. In the bottom TRF plot of each panel, the colored parts represent the electrodes within the significant cluster and the integration window over which they are significant.

#### S.1.4.3 Lesion volume in aphasia subgroups

A significant difference was found between the lesion volume in the mild aphasia group and the more severe aphasia group (F(1, 37)=14.307; p=0.0005; fig. S.10).

**Figure S.10:**
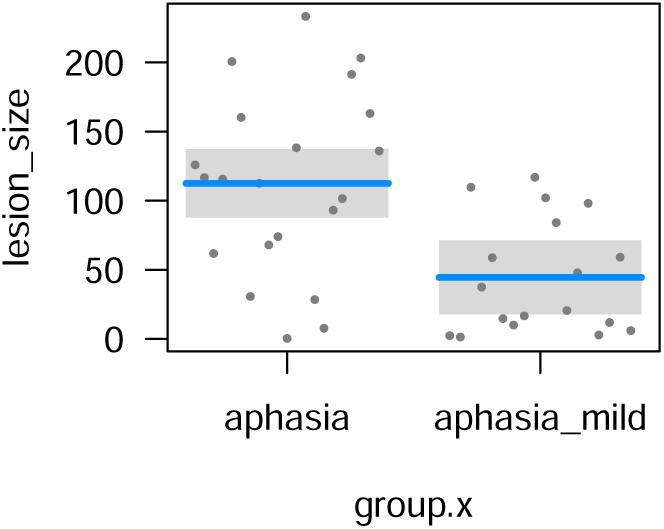
Lesion volume of the mild aphasia and more severe aphasia group. The lesion volume is significantly larger in the more severe aphasia group as compared to the mild aphasia group.

### S.1.5 Are IWA with a stroke origin in the posterior cerebral artery driving the posterior clusters?

Sub-010, sub-017 and sub-019 had a stroke originating in the posterior cerebral artery, which can affect occipital cortex, but also inferior/ middle temporal gyrus and parts of the parietal cortex. We were wondering whether the posterior clusters that we observe in the cluster-based permutation tests (figures 4 and 5) may be driven by the 3 IWA who had a lesion originating from the posterior cerebral artery. Therefore, we repeated the cluster-based permutation tests for the acoustic model prediction accuracy and envelope TRFs without sub-010, sub-017 and sub-019.

In figure S.11, we show the results. The clusters stay very similar as to when we conduct the tests with all 39 IWA, and the posterior clusters remain present. The 3 left-out subjects do thus not drive the posterior clusters that we observe.

When we conduct the group comparison on the electrode-average (with covariates, including lesion size; see section 2.4.1), the group difference also stays significant when the suggested 3 IWA are excluded (F=6.198; p=0.016).

This supplementary analysis shows that the posterior clusters are not driven by the lesion site of the 3 IWA who had a lesion originating from the posterior cerebral artery.

**Figure S.11:**
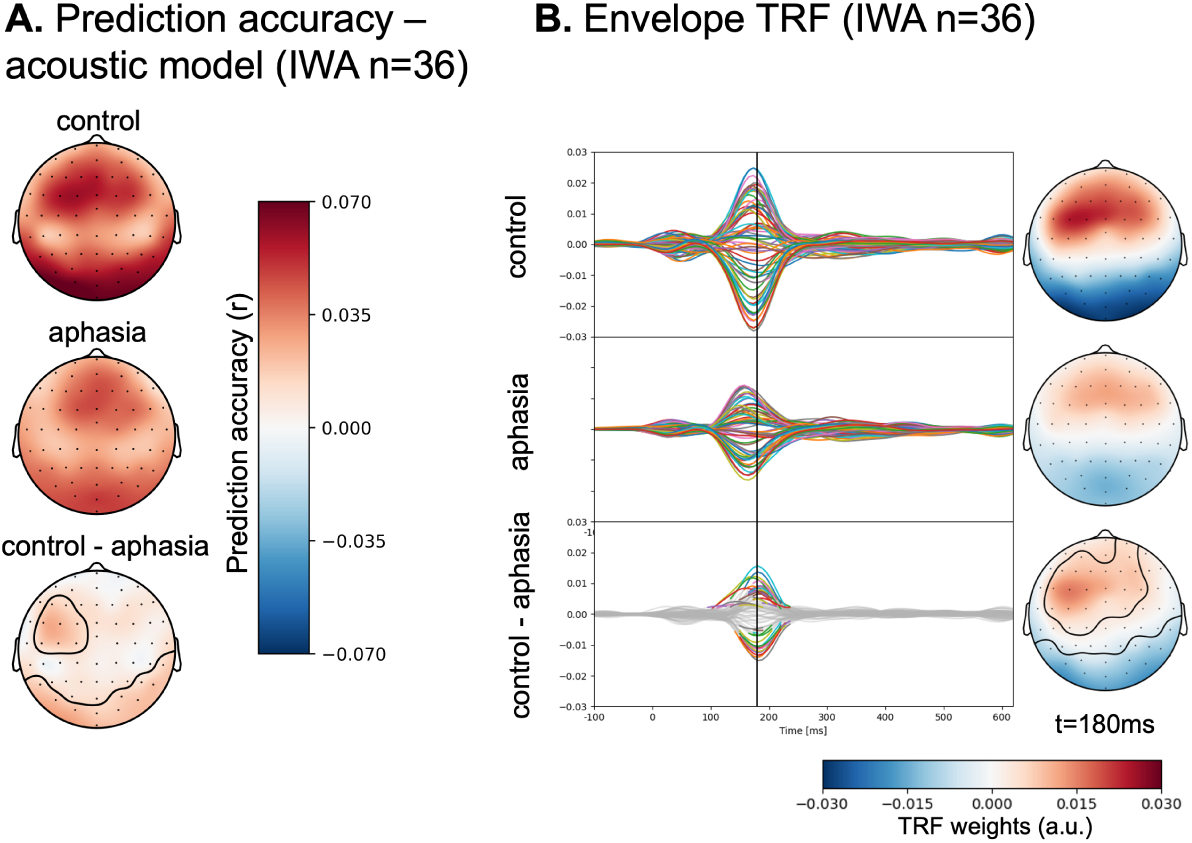
The patterns resulting from the cluster-based permutations when the 3 IWA with lesions resulting from PCA damage are excluded are similar to the pattern of results when the whole aphasia group is compared to the control group. In the encircled clusters, the aphasia group had significantly lower amplitudes than the control group.

## References

1. Aerts, A., van Mierlo, P., Hartsuiker, R. J., Santens, P., and De Letter, M. (2015). Neurophysiological sensitivity for impaired phonological processing in the acute stage of aphasia. Brain and Language, 149:84–96.

2. Aiken, S. J. and Picton, T. W. (2008). Human cortical responses to the speech envelope. Ear and Hearing, 29(2):139–157.

3. Anwyl-Irvine, A. L., Massonnié, J., Flitton, A., Kirkham, N., and Evershed, J. K. (2020). Gorilla in our midst: An online behavioral experiment builder. Behavior Research Methods, 52(1):388–407.

4. Becker, F. and Reinvang, I. (2007). Successful syllable detection in aphasia despite processing impairments as revealed by event-related potentials. Behavioral and Brain Functions, 3:1–16.

5. Biesmans, W., Das, N., Francart, T., and Bertrand, A. (2017). Auditory-inspired speech envelope extraction methods for improved eeg-based auditory attention detection in a cocktail party scenario. IEEE Transactions on Neural Systems and Rehabilitation Engineering, 25(5):402–412.

6. Brady, M. C. (2022). Dosage, intensity, and frequency of language therapy for aphasia: A systematic review-based, individual participant data network meta-analysis. Stroke, 29:956–967.

7. Brodbeck, C. (2020). Eelbrain 0.32. 10.5281/zenodo.3923991.

8. Brodbeck, C., Das, P., Gillis, M., Kulasingham, J. P., Bhattasali, S., Gaston, P., Resnik, P., and Simon, J. Z. (2021). Eelbrain: A python toolkit for time-continuous analysis with temporal response functions. BioRxiv.

9. Brodbeck, C., Presacco, A., and Simon, J. Z. (2018). Neural source dynamics of brain responses to continuous stimuli: Speech processing from acoustics to comprehension. NeuroImage, 172:162–174.

10. Broderick, M. P., Anderson, A. J., Di Liberto, G. M., Crosse, M. J., and Lalor, E. C. (2018). Electrophysiological correlates of semantic dissimilarity reflect the comprehension of natural, narrative speech. Current Biology, 28:803–809.e3.

11. Broderick, M. P., Anderson, A. J., and Lalor, E. C. (2019). Semantic context enhances the early auditory encoding of natural speech. Journal of Neuroscience, 39(38):7564–7575.

12. Broderick, M. P., Di Liberto, G. P., Anderson, A. J., Rofes, A., and Lalor, E. C. (2021). Dissociable electrophysiological measures of natural language processing reveal differences in comprehension strategy in healthy ageing. Scientific reports, 11(4963):1–12.

13. Brumm, K. P., Perthen, J. E., Liu, T. T., Haist, F., Ayalon, L., and Love, T. (2010). An arterial spin labeling investigation of cerebral blood flow deficits in chronic stroke survivors. NeuroImage, 51(2010):995–1005.

14. Cassidy, J. M., Wodeyar, A., Wu, J., Kaur, K., Masuda, A. K., Srinivasan, R., and Cramer, S. C. (2020). Low-frequency oscillations are a biomarker of injury and recovery after stroke. Stroke, 51(5):1442–1450.

15. Chang, C. T., Lee, C. Y., Chou, C. J., Fuh, J. L., and Wu, H. C. (2016). Predictability effect on N400 reflects the severity of reading comprehension deficits in aphasia. Neuropsychologia, 81:117–128.

16. Cocquyt, E.-M., Vandewiele, M., Bonnarens, C., Santens, P., and De Letter, M. (2020). The sensitivity of event-related potentials/fields to logopedic interventions in patients with stroke-related aphasia. Acta Neurologica Belgica, 120(4):2240–2993.

17. Cohen, R., Abboud, S., and Arad, M. (2015). Monitoring brain damage using bioimpedance technique in a 3d numerical model of the head. Medical Engineering and Physics, 37(5):453–459.

18. Cordella, C., Munsell, M., Godlove, J., Anantha, V., Advani, M., and Kiran, S. (2022). Dosage frequency effects on treatment outcomes following self-managed digital therapy: Retrospective cohort study. Journal of Medical Internet Research, 24(7):1–14.

19. David, S. V., Mesgarani, N., and Shamma, S. A. (2007). Estimating sparse spectro-temporal receptive fields with natural stimuli. Network: Computation in Neural Systems, 18:191–212.

20. De Clercq, P., Kries, J., Mehraram, R., Vanthornhout, J., Francart, T., and Vandermosten, M. (2023). Detecting post-stroke aphasia using eeg-based neural envelope tracking of natural speech. *medrxiv*.

21. Di Liberto, G., Peter, V., Kalashnikova, M., Goswami, U., Burnham, D., and Lalor, E. (2018). Atypical cortical entrainment to speech in the right hemisphere underpins phonemic deficits in dyslexia. NeuroImage, 175:70–79.

22. Di Liberto, G. M. and Lalor, E. C. (2017). Indexing cortical entrainment to natural speech at the phonemic level: Methodological considerations for applied research. Hearing Research, 348:70–77.

23. Di Liberto, G. M., O’Sullivan, J. A., and Lalor, E. C. (2015). Low frequency cortical entrainment to speech reflects phoneme level processing. Current Biology, 25:2457–2465.

24. Dial, H. R., Gnanateja, G. N., Tessmer, R. S., Gorno-Tempini, M. L., Chandrasekaran, B., and Henry, M. L. (2021). Cortical tracking of the speech envelope in logopenic variant primary progressive aphasia. Frontiers in Human Neuroscience, 14(597694):1–9.

25. Ding, N. and Simon, J. Z. (2012). Neural coding of continuous speech in auditory cortex during monaural and dichotic listening. Journal of Neurophysiology, 107:78–89.

26. Duchateau, J., Kong, Y. O., Cleuren, L., Latacz, L., Roelens, J., Samir, A., Demuynck, K., Ghesquière, P., Verhelst, W., and Van hamme, H. (2009). Developing a reading tutor: Design and evaluation of dedicated speech recognition and synthesis modules. Speech Communication, 51(10):985–994.

27. El Hachioui, H., Visch-Brink, E. G., Lingsma, H. F., Van De Sandt-Koenderman, M. W., Dippel, D. W., Koudstaal, P. J., and Middelkoop, H. A. (2014). Nonlinguistic cognitive impairment in poststroke aphasia: A prospective study. Neurorehabilitation and Neural Repair, 28(3):273–281.

28. Engelter, S. T., Gostynski, M., Papa, S., Frei, M., Born, C., Ajdacic-Gross, V., Gutzwiller, F., and Lyrer, P. A. (2006). Epidemiology of aphasia attributable to first ischemic stroke: Incidence, severity, fluency, etiology, and thrombolysis. Stroke, 37(6):1379–1384.

29. Federmeier, K. D., McLennan, D. B., de Ochoa, E., and Kutas, M. (2002). The impact of semantic memory organization and sentence context information on spoken language processing by younger and older adults: An ERP study. Psychophysiology, 39(2):133–146.

30. Flamand-Roze, C., Falissard, B., Roze, E., Maintigneux, L., Beziz, J., Chacon, A., Join-Lambert, C., Adams, D., and Denier, C. (2011). Validation of a new language screening tool for patients with acute stroke. Stroke, 42(5):1224–1229.

31. Fló, A., Brusini, P., Macagno, F., Nespor, M., Mehler, J., and Ferry, A. L. (2019). Newborns are sensitive to multiple cues for word segmentation in continuous speech. Developmental Science, 22(4):1–16.

32. Fonseca, J., Raposo, A., and Martins, I. P. (2018). Cognitive functioning in chronic post-stroke aphasia. Applied Neuropsychology:Adult, 26(4):355–364.

33. Francart, T., Wieringen, A. V., and Wouters, J. (2008). Apex 3: a multi-purpose test platform for auditory psychophysical experiments. J Neurosci Methods, 172:283–293.

34. Fridriksson, J., Ouden, D. B. D., Hillis, A. E., Hickok, G., Rorden, C., Basilakos, A., Yourganov, G., and Bonilha, L. (2018). Anatomy of aphasia revisited. Brain, 141:848–862.

35. Gaspers, J., Cimiano, P., Rohlfing, K., and Wrede, B. (2017). Constructing a language from scratch: Combining bottom–up and top–down learning processes in a computational model of language acquisition. IEEE Transactions on Cognitive and Developmental Systems, 9(2):183–196.

36. Gillis, M., Van Canneyt, J., Francart, T., and Vanthornhout, J. (2022). Neural tracking as a diagnostic tool to assess the auditory pathway. Hearing Research, 426(108607):1–14.

37. Gillis, M., Vanthornhout, J., Simon, J. Z., Francart, T., and Brodbeck, C. (2021). Neural markers of speech comprehension: measuring eeg tracking of linguistic speech representations, controlling the speech acoustics. Journal of Neuroscience, 41(50):10316–10329.

38. Gillis and Kries, Vandermosten, M., and Francart, T. (2023). Neural tracking of linguistic and acoustic speech representations decreases with advancing age. NeuroImage, 267(119841):1–16.

39. Hamilton, L. S. and Huth, A. G. (2018). The revolution will not be controlled: natural stimuli in speech neuroscience. *Language*, Cognition and Neuroscience, 35(5):573–582.

40. Harris, K. C. (2020). The Aging Auditory System: Electrophysiology. In Helfer, K. S., Bartlett, E. L., Popper, A. N., and Fay, R. R., editors, Aging and Hearing: Causes and Consequences, pages 117–141. Springer International Publishing, Cham.

41. Hickok, G. and Poeppel, D. (2007). The cortical organization of speech processing. Nature Reviews Neuroscience, 8(May):393–402.

42. Hillyard, S. and Kutas, M. (1984). Brain potentials during reading reflect word expectancy and semantic association. Nature, 307(5947):161–163.

43. Holdgraf, C. R., Rieger, J. W., Micheli, C., Martin, S., Knight, R. T., and Theunissen, F. E. (2017). Encoding and decoding models incognitive electrophysiology. Frontiers in Systems Neuroscience, 11(61):1– 24.

44. Huygelier, H., Schraepen, B., Demeyere, N., and Gillebert, C. R. (2019). The Dutch version of the Oxford Cognitive Screen (OCS-NL): normative data and their association with age and socio-economic status. *Aging*, Neuropsychology, and Cognition, 27(5):765–786.

45. Ilvonen, T., Kujala, T., Kozou, H., Kiesiläinen, A., Salonen, O., Alku, P., and Näätänen, R. (2004). The processing of speech and non-speech sounds in aphasic patients as reflected by the mismatch negativity (MMN). Neuroscience Letters, 366(3):235–240.

46. Ilvonen, T.-M., Kujala, T., Tervaniemi, M., Salonen, O., Näätänen, R., and Pekkonen, E. (2001). The processing of sound duration after left hemisphere stroke: Event-related potential and behavioral evidence. Psychophysiology, 38:622–628.

47. Kandylaki, K. D. and Bornkessel-Schlesewsky, I. (2019). From story comprehension to the neurobiology of language. Language, Cognition and Neuroscience, 34(4):405–410.

48. Kawohl, W., Bunse, S., Willmes, K., Hoffrogge, A., Buchner, H., and Huber, W. (2010). Semantic eventrelated potential components reflect severity of comprehension deficits in aphasia. Neurorehabilitation and Neural Repair, 24(3):282–289.

49. Keuleers, E., Brysbaert, M., and New, B. (2010). Subtlex-nl: A new measure for dutch word frequency based on film subtitles. Behavior research methods, 42(3):643–650.

50. Khachatryan, E., De Letter, M., Vanhoof, G., Goeleven, A., and Van Hulle, M. M. (2017). Sentence context prevails over word association in aphasia patients with spared comprehension: Evidence from N400 event-related potential. Frontiers in Human Neuroscience, 10(684):1–15.

51. Kielar, A., Meltzer-Asscher, A., and Thompson, C. K. (2012). Electrophysiological responses to argument structure violations in healthy adults and individuals with agrammatic aphasia. Neuropsychologia, 50(14):3320–3337.

52. Kries, J., Clercq, P. D., Lemmens, R., Francart, T., and Vandermosten, M. (2023). Acoustic and phonemic processing are impaired in individuals with aphasia. Scientific Reports, 13.

53. Kuperberg, G. R. and Jaeger, T. F. (2016). What do we mean by prediction in language comprehension? Language, Cognition and Neuroscience, 31(1):32–59.

54. Kutas, M. and Federmeier, K. D. (2011). Thirty Years and Counting: Finding Meaning in the N400 Component of the Event-Related Brain Potential (ERP). Annual Review of Psychology, 62(1):621– 647.

55. Lalor, E. C. and Foxe, J. J. (2010). Neural responses to uninterrupted natural speech can be extracted with precise temporal resolution. European Journal of Neuroscience, 31:189–193.

56. Le, D., Licata, K., and Mower Provost, E. (2018). Automatic quantitative analysis of spontaneous aphasic speech. Speech Communication, 100:1–12.

57. Lesenfants, D. and Francart, T. (2020). The interplay of top-down focal attention and the cortical tracking of speech. Scientific Reports, 10(6922):1–10.

58. Lice, K. and Palmović, M. (2017). Semantic categorization in aphasic patients with impaired language comprehension: An event–related potentials study. Suvremena lingvistika, 43(84):135–155.

59. Lopopolo, A. and Rabovsky, M. (2022). Tracking lexical and semantic prediction error underlying the n400 using artificial neural network models of sentence processing. biorxiv.

60. Makov, S., Sharon, O., Ding, N., Ben-Shachar, M., Nir, Y., and Zion Golumbic, E. (2017). Sleep disrupts high-level speech parsing despite significant basic auditory processing. Journal of Neuroscience, 37:7772–7781.

61. Martin, B. A., Tremblay, K. L., and Korczak, P. (2008). Speech Evoked Potentials: From the Laboratory to the Clinic. Ear and Hearing, 29:285–313.

62. MATLAB (2016). version 9.1.0.441655 (R2016b). The MathWorks Inc., Natick, Massachusetts.

63. Mesgarani, N., Cheung, C., Johnson, K., and Chang, E. F. (2014). Phonetic feature encoding in human superior temporal gyrus. Science, 343(2):1006–1010.

64. Mesik, J., Ray, L., and Wojtczak, M. (2021). Effects of Age on Cortical Tracking of Word-Level Features of Continuous Competing Speech. Frontiers in Neuroscience, 15(4):1–21.

65. Michaelov, J. A. and Bergen, B. K. (2020). How well does surprisal explain n400 amplitude under different experimental conditions? In Proceedings of the 24th Conference on Computational Natural Language Learning, pages 652–663, online. Association for Computational Linguistics.

66. Mitra, V., Nam, H., Espy-Wilson, C. Y., Saltzman, E., and Goldstein, L. (2010). Retrieving tract variables from acoustics: A comparison of different machine learning strategies. IEEE J Sel Top Signal Process, 4:1027–1045.

67. National Aphasia Association (Accessed in August 2022). Aphasia fact sheet. NAA online publication.

68. Nieuwland, M. S., Barr, D. J., Bartolozzi, F., Busch-Moreno, S., Darley, E., Donaldson, D. I., Ferguson, H. J., Fu, X., Heyselaar, E., Huettig, F., Husband, E. M., Ito, A., Kazanina, N., Kogan, V., Kohút, Z., Kulakova, E., Mézière, D., Politzer-Ahles, S., Rousselet, G., Rueschemeyer, S. A., Segaert, K., Tuomainen, J., and Von Grebmer Zu Wolfsthurn, S. (2020). Dissociable effects of prediction and integration during language comprehension: Evidence from a largescale study using brain potentials. Philosophical Transactions of the Royal Society B: Biological Sciences, 375(1791):1–9.

69. Ofek, E., Purdy, S. C., Ali, G., Webster, T., Gharahdaghi, N., and McCann, C. M. (2013). Processing of emotional words after stroke: An electrophysiological study. Clinical Neurophysiology, 124:1771–1778.

70. Oganian, Y. and Chang, E. F. (2019). A speech envelope landmark for syllable encoding in human superior temporal gyrus. Science Advances, 5(11):1–13.

71. Oldfield, R. (1971). The assessment and analysis of handedness: the edinburgh inventory. Neuropsychologia, 9(1):97–113.

72. Park, W., Kwon, G. H., Kim, Y. H., Lee, J. H., and Kim, L. (2016). Eeg response varies with lesion location in patients with chronic stroke. Journal of NeuroEngineering and Rehabilitation, 13(21):1–10.

73. Pasley, B. N. and Knight, R. T. (2013). Decoding speech for understanding and treating aphasia. Progress in Brain Research, 207:435–456.

74. Peelle, J. E., Gross, J., and Davis, M. H. (2013). Phase-locked responses to speech in human auditory cortex are enhanced during comprehension. Cerebral Cortex, 23(6):1378–1387.

75. Pettigrew, C. M., Murdoch, B. E., Kei, J., Ponton, C. W., Alku, P., and Chenery, H. J. (2005). The mismatch negativity (MMN) response to complex tones’ and spoken words in individuals with aphasia. Aphasiology, 19(2):131–163.

76. Piastra, M. C., Oostenveld, R., Schoffelen, J. M., and Piai, V. (2022). Estimating the influence of stroke lesions on meg source reconstruction. NeuroImage, 260:1–13.

77. Prinsloo, K. D. and Lalor, E. C. (2022). General auditory and speech-specific contributions to cortical envelope tracking revealed using auditory chimeras. Journal of Neuroscience, 42(41):7782–7798.

78. Pulvermüller, F., Mohr, B., and Lutzenberger, W. (2004). Neurophysiological correlates of word and pseudo-word processing in well-recovered aphasics and patients with right-hemispheric stroke. Psychophysiology, 41(4):584–591.

79. R Core Team (2017). R: A Language and Environment for Statistical Computing. R Foundation for Statistical Computing, Vienna, Austria.

80. Rabiller, G., He, J.-W., Nishijima, Y., Wong, A., and Liu, J. (2015). Perturbation of brain oscillations after ischemic stroke: A potential biomarker for post-stroke function and therapy. Int. J. Mol. Sci, 16:25605–25640.

81. Räling, R., Schröder, A., and Wartenburger, I. (2016). The origins of age of acquisition and typicality effects: Semantic processing in aphasia and the ageing brain. Neuropsychologia, 86:80–92.

82. Robson, H., Pilkington, E., Evans, L., DeLuca, V., and Keidel, J. L. (2017). Phonological and semantic processing during comprehension in Wernicke’s aphasia: An N400 and Phonological Mapping Negativity Study. Neuropsychologia, 100:144–154.

83. Rohde, A., Worrall, L., Godecke, E., O’Halloran, R., Farrell, A., and Massey, M. (2018). Diagnosis of aphasia in stroke populations: A systematic review of language tests. PLoS ONE, 13(3):1–17.

84. Salinet, A. S. M., Panerai, R. B., Robinson, T. G., and Salinet, A. (2014). The longitudinal evolution of cerebral blood flow regulation after acute ischaemic stroke. Cerebrovasc Dis Extra, 4:186–197.

85. Schevenels, K., Gerrits, R., Lemmens, R., De Smedt, B., Zink, I., and Vandermosten, M. (2022). Early white matter connectivity and plasticity in post stroke aphasia recovery. NeuroImage: Clinical, 36:1–12.

86. Schevenels, K., Price, C. J., Zink, I., De Smedt, B., and Vandermosten, M. (2020). A review on treatmentrelated brain changes in aphasia. Neurobiology of Language, 1(4):402–433.

87. Shannon, R. V., Zeng, F.-G., Kamath, V., Wygonski, J., and Ekelid, M. (1995). Speech recognition with primarily temporal cues. Science, 270(5234):303–304.

88. Sheppard, S. M., Love, T., Midgley, K. J., Holcomb, P. J., and Shapiro, L. P. (2017). Electrophysiology of prosodic and lexical-semantic processing during sentence comprehension in aphasia. Neuropsychologia, 107:9–24.

89. Shuai, L., Gong, T., Giuliano, R., and Gu, W. (2014). Temporal relation between top-down and bottomup processing in lexical tone perception. Frontiers in Behavioral Neuroscience, 8(97):1–16.

90. Silkes, J. A. P. and Anjum, J. (2021). The role and use of event-related potentials in aphasia: A scoping review. Brain and Language, 219:1–16.

91. Smith, L. and Yu, C. (2008). Infants rapidly learn word-referent mappings via cross-situational statistics. Cognition, 106(3):1558–1568.

92. Somers, B., Francart, T., and Bertrand, A. (2018). A generic eeg artifact removal algorithm based on the multi-channel wiener filter. Journal of Neural Engineering, 15(3):1–13.

93. Spreng, R. N. and Turner, G. R. (2019). The Shifting Architecture of Cognition and Brain Function in Older Adulthood. Perspectives on Psychological Science, 14(4):523–542.

94. Suanda, S. H., Mugwanya, N., and Namy, L. L. (2014). Cross-situational statistical word learning in young children. Journal of Experimental Child Psychology, 126:395–411.

95. Thiessen, E. and Erickson, L. (2013). Discovering words in fluent speech: The contribution of two kinds of statistical information. Frontiers in Psychology, 3:1–10.

96. Tremblay, P. and Dick, A. S. (2016). Broca and wernicke are dead, or moving past the classic model of language neurobiology. Brain and Language, 162:60–71.

97. Van Ewijk, E., Dijkhuis, L., Hofs-Van Kats, M., Hendrickx-Jessurun, M., Wijngaarden, M., and De Hilster, C. (2020). Nederlandse Benoem Test. Bohn Stafleu Van Loghum, Houten, NL.

98. Van Rossum, G. and Drake Jr, F. L. (1995). Python reference manual. Centrum voor Wiskunde en Informatica Amsterdam.

99. Vanthornhout, J., Decruy, L., Wouters, J., Simon, J. Z., and Francart, T. (2018). Speech intelligibility predicted from neural entrainment of the speech envelope. JARO – Journal of the Association for Research in Otolaryngology, 19(2):181–191.

100. Verwimp, L., Van hamme, H., and Wambacq, P. (2019). Tf-lm: Tensorflow-based language modeling toolkit. In http://www.lrec-conf.org/proceedings/lrec2018/index.html, pages 2968–2973. Proceedings LREC.

101. Visch-Brink, E., Van de Sandt-Koenderman, M., and El Hachioui, H. (2010). ScreeLing. Houten: Bohn Stafleu Van Loghum.

102. Vorwerk, J., Cho, J. H., Rampp, S., Hamer, H., Knösche, T. R., and Wolters, C. H. (2014). A guideline for head volume conductor modeling in eeg and meg. NeuroImage, 100:590–607.

103. Weissbart, H., Kandylaki, K. D., and Reichenbach, T. (2019). Cortical tracking of surprisal during continuous speech comprehension. Journal of Cognitive Neuroscience, 32(1):155–166.

104. Wilson, S. M., Entrup, J. L., Schneck, S. M., Onuscheck, C. F., Levy, D. F., Rahman, M., Willey, E., Casilio, M., Yen, M., Brito, A. C., Kam, W., Davis, L. T., Riesthal, M. D., and Kirshner, H. S. (2023). Recovery from aphasia in the first year after stroke. Brain, 146:146–1021.

105. Wlotko, E. W., Lee, C. L., and Federmeier, K. D. (2010). Language of the Aging Brain: Event-Related Potential Studies of Comprehension in Older Adults. Linguistics and Language Compass, 4(8):623–638.

106. Xu, L. and Pfingst, B. E. (2008). Spectral and temporal cues for speech recognition: Implications for auditory prostheses. Hearing Research, 242(1-2):132–140.

107. Zbesko, J. C., Nguyen, T.-V. V., Yang, T., Frye, J. B., Hussain, O., Hayes, M., Chung, A., Day, W. A., Stepanovic, K., Krumberger, M., Mona, J., Longo, F. M., and Doyle, K. P. (2018). Glial scars are permeable to the neurotoxic environment of chronic stroke infarcts. Neurobiol Dis, 112:63–78.

108. Zeng, F. G., Nie, K., Stickney, G. S., Kong, Y. Y., Vongphoe, M., Bhargave, A., Wei, C., and Cao, K. (2005). Speech recognition with amplitude and frequency modulations. Proceedings of the National Academy of Sciences of the United States of America, 102(7):2293–2298.

